# Comparative analysis of Illumina and Ultima-Genomics sequencing for plasma cell-free small RNA profiling in pancreatic cancer

**DOI:** 10.64898/2026.06.21.733585

**Authors:** Amit Levon, Hadas Volkov, Rani Shlayem, Noam Shomron

**Affiliations:** Gray Faculty of Medical and Health Sciences, Tel Aviv University, Tel Aviv, Israel; Edmond J. Safra Center for Bioinformatics, Tel Aviv University, Tel Aviv, Israel; Sagol School of Neuroscience, Tel Aviv University, Tel Aviv, Israel

**Keywords:** Ultima-Genomics, Illumina, Cell-free RNA, MicroRNA, NGS, Bioinformatics

## Abstract

Plasma-derived cell-free small non-coding RNAs are promising non-invasive biomarkers for cancer detection and monitoring. However, variability in sequencing output limits standardization, and cross-platform performance for plasma small RNA profiling has not been systematically evaluated. Illumina short-read sequencing is the current standard, whereas the newcomer, Ultima-Genomics platform, has been less extensively studied for circulating small RNA in plasma. To directly compare platform performance, we sequenced plasma cell-free RNA from 39 patients with pancreatic cancer and 39 matched controls on both platforms. After filtering, Ultima-Genomics retained more mature microRNA reads, whereas Illumina achieved slightly higher enrichment efficiency and mapping rates. Despite these technical differences, both platforms produced concordant expression profiles, with strong cross-platform correlations for shared microRNAs and clear separation of cases and controls within each dataset. Differential expression analysis identified 14 significant microRNAs on both platforms with concordant directions of change, most of which are supported by pancreatic cancer databases. Pathway enrichment analysis highlighted signaling pathways implicated in pancreatic cancer, supporting the biological relevance of both shared and platform-specific signatures. These findings indicate that both Illumina and Ultima Genomics platforms are suitable for plasma small RNA profiling and capture biologically relevant signals in pancreatic cancer.

## Introduction

Pancreatic ductal adenocarcinoma (PDAC) is among the most lethal types of cancer, with a survival rate of around five years^1^. The majority of patients are diagnosed at advanced stages, with unresectable, locally advanced, or metastatic disease stages^2^. This poor prognosis is driven in large part by the lack of sensitive, non-invasive tools for early detection and for monitoring minimal residual disease (MRD). Conventional serological biomarkers (such as CA19-9) have limited sensitivity and specificity, especially in early-stage PDAC^3^. Therefore, liquid biopsy approaches that analyze tumor-derived material in blood have considerable promise for improving PDAC diagnosis and disease monitoring^4–6^.

Specifically, plasma-derived cell-free RNAs (cfRNAs), and especially small non-coding RNAs (cf-sRNAs), allow a dynamic, systemic and non-invasive view of tumor biology^5,7^. Beyond acting as passive markers, miRNAs actively regulate oncogenic pathways and drive metastasis across various cancers^8–10^. Multiple studies have reported PDAC-associated changes in circulating miRNA profiles^11,12^. The cf-sRNA panels can even differentiate patients with resectable PDAC from controls and enhance the performance of CA19-9 when used in combination with the marker^4^. These findings make cf-sRNAs attractive biomarker candidates for early detection and risk stratification in PDAC^13^.

However, transforming cfRNA signatures into robust clinical assays is difficult due to technical variability across RNA-seq workflows^14^. Systematic cross-study analyses of plasma cfRNA-seq data show that factors such as genomic DNA contamination, microbial sequences, and library-specific biases can dominate the observed transcriptomic variation, which can overshadow true biological differences between patient groups^15^. This means that sequencing platform choice and sequencing chemistry are not just practical considerations, but are key factors affecting data quality, biotype representation, and the reproducibility of cf-sRNA biomarker discovery.

To date, Illumina short-read sequencing has served as the industry standard for plasma cfRNA and cf-sRNA profiling, underpinning the majority of published cfRNA workflows and benchmarking efforts¹¹. Recently, the Ultima-Genomics (UG) platform, a technology based on mostly natural sequencing-by-synthesis (mnSBS) and an open-fluidics spinning-wafer architecture, was developed as a high-throughput, cost-efficient comparable alternative that can deliver ∼30× whole-genome coverage^16,17^. Studies have shown that UG sequencing results in gene expression and variant-calling performance are comparable to Illumina for bulk RNA-seq, single-cell RNA-seq, and deep whole-genome ctDNA profiling, while allowing deeper or broader sequencing within a fixed budget^18–20^. Nevertheless, systematic evaluations of UG for plasma-derived cf-sRNA are still lacking.

We address this gap by sequencing the same plasma-derived cf-sRNA cohort from PDAC patients and matched controls on both Illumina and UG platforms, using an identical small-RNA library preparation workflow. We build on previous work that highlighted the dominant impact of technical factors on cfRNA-seq data, and perform a direct, head-to-head comparison of sequencing performance and data characteristics for cf-sRNA on these two platforms in a real-world PDAC liquid biopsy context. To our knowledge, this is the first study to sequence the same plasma-derived cfRNA cohort on both Illumina and UG platforms, enabling a direct technical comparison of sequencing performance and data characteristics in this clinically important analytical setting.

## Results

### Sequencing, quality control and alignment

Raw FASTQ files from both platforms underwent initial quality control (QC) and adapter/length filtering (18–40 nt), resulting in the ‘Total Short cfRNA’ dataset for downstream analysis. These were then processed, producing mature miRNA reads after contamination depletion and miRBase alignment (see Methods). All subsequent QC metrics compare these two datasets (“Total Short cfRNA” vs “Mature miRNA”). We first assess the baseline performance of Ultima UG 100 (flow-based non-terminating chemistry) and Illumina NovaSeq 6000 (sequencing-by-synthesis; SBS).

Ultima UG 100 generated substantially more raw reads per sample than Illumina NovaSeq 6000, and this difference persisted after adapter trimming (mean 21.5M reads, std 7.8M, vs 4.2M reads, std 1.6M; Fig. S1a and S1b, Table S1). After decontamination and alignment to mature miRNAs, UG retained ∼3.8-fold more mature miRNA reads per sample than Illumina (mean 56K, std 47K, vs 15K, std 13K; Fig. S1a and S1c, Table S1). However, mature miRNA reads represented a slightly smaller fraction of total platform yield for UG than for Illumina (0.0033%, std 0.0028%, vs 0.0045%, std 0.0039%; Fig. 1c), indicating modestly higher enrichment efficiency for Illumina.

**Fig. 1:**
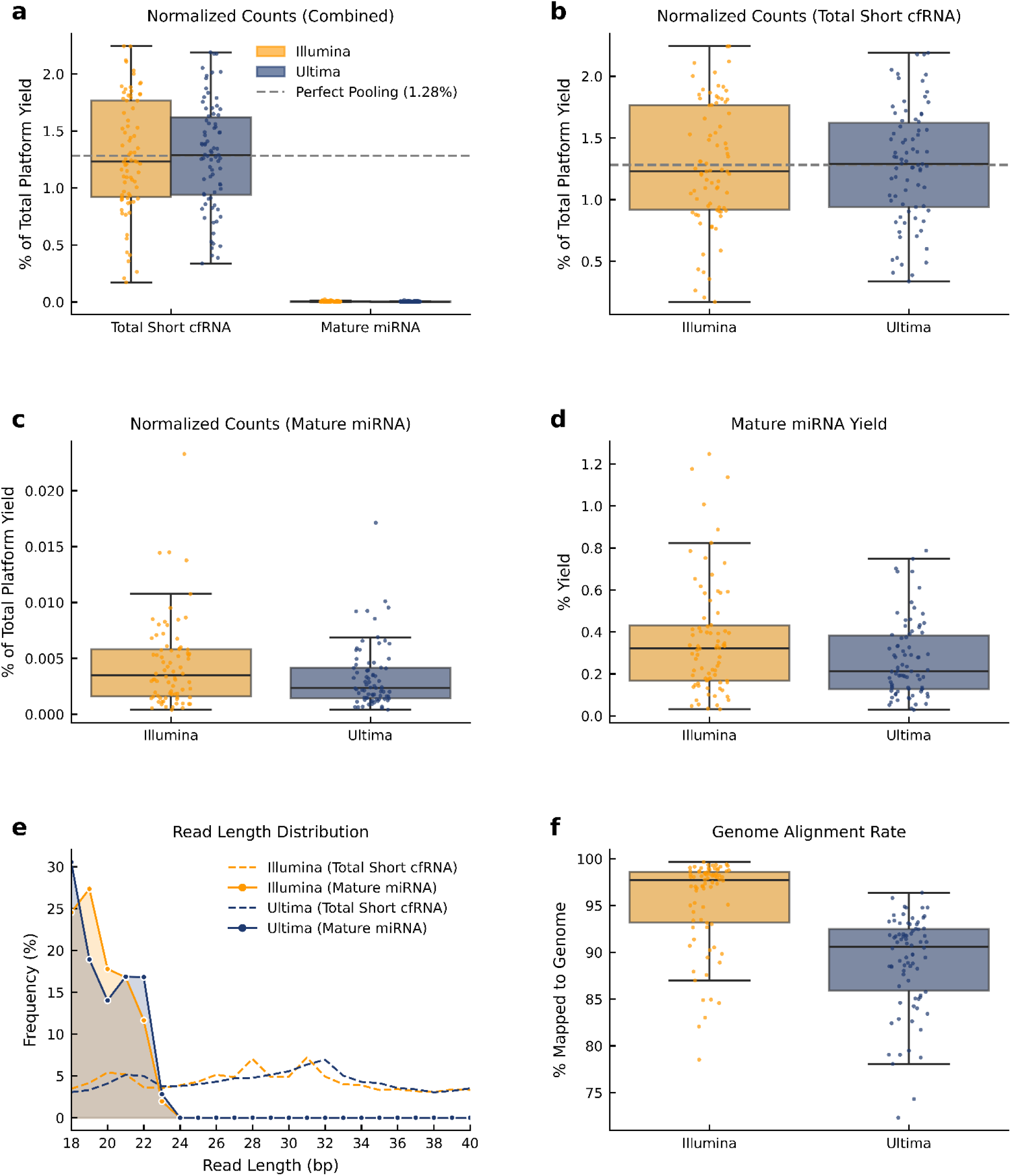
General sequencing and alignment comparisons between Illumina (orange) and UG (navy). **a** Combined per-library normalized read counts (Sample Reads / Total Platform Reads × 100) evaluated directly after post-trimming/QC (Total Short cfRNA) and after mature miRNA alignment. Boxplots with jitter points show medians, interquartile ranges, and individual library distributions (n=78 per platform). The horizontal gray dashed line represents theoretical perfect pooling equality (1.28%), the baseline where all 78 multiplexed samples are equally distributed within the sequencing run. **b** Isolated comparison of the normalized read counts post-trimming/QC (Total Short cfRNA). **c** Isolated comparison of the normalized read counts after mature miRNA alignment. **d** Mature miRNA yield - % of post-adapter trimming reads successfully aligning to mature miRNAs (percentage of post-trimming total short cfRNA reads aligning to mature miRNAs), by platform. **e** Read length distributions before and after miRNA-specific filtering. Mean read length frequency (% of reads) is shown across 18-40nt for Illumina and UG libraries (n=78 each). Dashed lines represent the baseline post-adapter trimming/QC (Total Short cfRNA) distributions. Solid lines with markers represent distributions following sequential decontamination filtering against cfDNA, ncRNA, rRNA, tRNA, and piRNA reference databases, keeping only reads mapping to miRBase mature miRNA loci. Shaded regions highlight the mature-miRNA-enriched size range, visually representing the canonical 18-24nt miRNA size enrichment. **f** Per-library % reads mapped to GRCh38 genome after contamination filtering. Boxplots with jitter points (n=78/platform). High rates validate alignment quality across platforms.

Mature miRNA alignment yield was low across platforms, as expected for plasma cf-sRNA, with Illumina averaging ∼0.36% (std ∼0.27%) and UG ∼0.27% (std ∼0.18%) (Fig. 1d). Thus, the UG libraries underwent more filtering during processing. This ∼25% higher relative yield for Illumina reflects slightly better target enrichment despite lower absolute depth. This is consistent with SBS chemistry’s lower off-target artifacts compared to UG’s flow cell-based approach. Despite this higher loss, the final usable read depth for UG remains higher than Illumina (Fig. S1c).

General total short cfRNA read length distributions show broad, multi-modal profiles (Fig. 1e), as expected from plasma-derived short nucleic acid (NA) fragments. Both platforms show an early peak at ∼19-22 nt corresponding to the miRNA-length window, but diverge at longer lengths: Illumina total short cfRNA reads demonstrate secondary peaks near 28 and 31 nt (∼7%), while UG total short cfRNA reads exhibit a flatter, more uniform distribution across 24-40 nt (∼3-7%) with rising modest elevation near 32 nt. These longer fragments are likely caused by the abundant presence of co-sequenced non-miRNAs, common in plasma total NA like native short cfDNA^21^ and structured RNA fragments (tRNA halves, Y-RNAs, and 5.8S rRNA fragments)^22^. Sequential mapping against non-coding RNA reference databases confirms their identity and abundance: cDNA (cfDNA) represented the largest filtered fraction (∼34-37% of reads), followed by ncRNA (∼12-15%), and rRNA (∼8-10%) for both platforms (Fig. S2a; Table S2). Following this filtering step, read length distributions narrow sharply to canonical miRNA lengths below 24 nt (Fig. 1e), confirming isolation of the mature miRNA fraction and the non-miRNA origin of the longer fragment signal.

Mature miRNA distributions narrowed sharply to the canonical under 22 nt lengths. Canonical mature miRNAs typically span 19-25 nt (averaging at 22 nt; Fig. S3a), yet we observed an elevated left-shoulder with distinct local peaks at 18-19 nt in both platforms (Fig. 1e). The subdued 21-23 nt values and enrichment of shorter sequences may be caused by 3’ extracellular processing and exonuclease trimming in of canonical miRNAs into shorter isomiRs in circulation in the plasma^23,24^. Another reason may be the high relative abundance of specific circulating miRNAs (such as miR-486-5p and miR-1246) whose most stable, predominant circulating isoforms are naturally 18-19 nt in length^25–27^ (Fig. S3b and S3c). Importantly, this length heterogeneity does not affect DESeq2 differential expression analysis, as isomiRs are collapsed to their parent miRNA locus during count matrix construction, and DESeq2 models total per-locus counts^28,29^.

Genome alignment rates were high and platform-comparable (Fig. 1f), with Illumina averaging 95% ± 5% mapped reads vs UG 89% ± 5% (medians 98% vs 91%; n=78 libraries each) and overlapping distributions indicating robust short-read mappability post-filtering. The modest but consistent mapping advantage of Illumina (mean 95.3% vs. 88.9%) is consistent with known differences in platform-specific error profiles. UG’s flow-based chemistry produces a higher proportion of indel-type errors compared to Illumina’s substitution-dominant SBS chemistry^16,30,31^, and for very short reads (18-24 nt), single-base insertions or deletions are more likely to prevent alignment than substitution errors. This difference in mapping rate did not prevent high-quality expression profiling, which is consistent with previous benchmarking of the UG 100 against Illumina for RNA-seq applications^20^. Additional QC metrics are shown in Figures S4-S6.

### sRNA types and miRNA statistics

Looking specifically at the mapped reads reveals differences in RNA types (Fig. 2a). Among genome-aligned reads, the majority remained unassigned/unknown after decontamination, ∼81% of Illumina (std ∼11%) and ∼88% of UG reads (std 8%). This is generally consistent with the fragmented and heterogeneous nature of plasma cf-sRNA, in which a large proportion of circulating small RNA reads do not overlap (or cannot be confidently assigned) to the standard biotype reference databases^32,33^. Extracellular/cell-free RNA libraries contain diverse short fragments from many RNA sources beyond canonical miRNA/tRNA/rRNA such as introns, lncRNAs, Y RNAs, sn/snoRNAs, misc RNAs, repeats, etc^34^. For example, tRNA gene catalogs are often incomplete/uncertain, mature tRNAs carry many chemical modifications that cause mismatches/deletions during reverse transcription, which makes “standard” mapping/assignment harder^35^. Therefore, the large unknown fraction reflects both extracellular RNA fragment diversity and the conservative annotation we used.

**Fig. 2:**
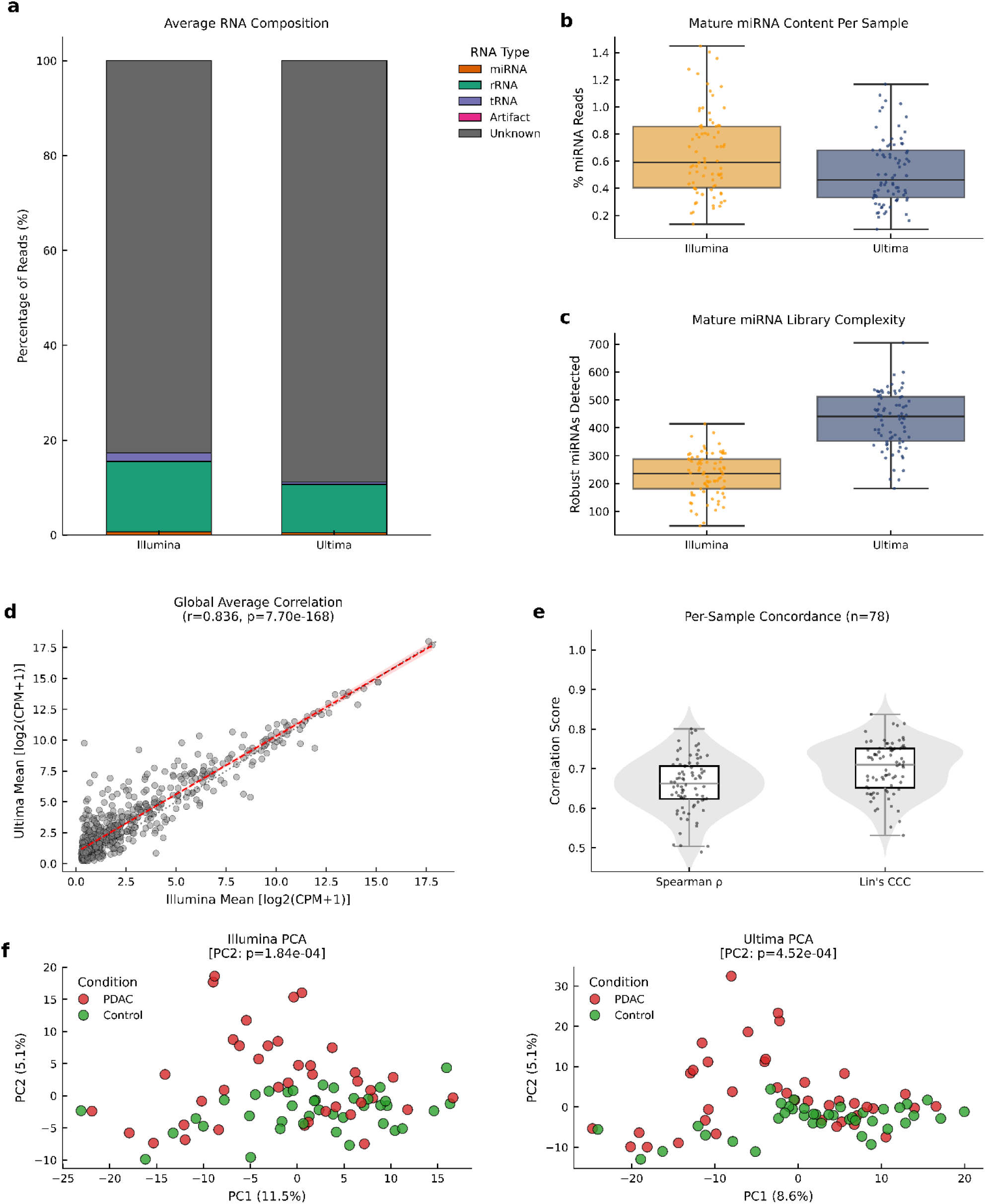
RNA composition, library complexity, and miRNA expression profiles across Illumina and UG platforms. **a** Mean RNA biotype composition (% of genome-mapped reads). Stacked bars show major RNA categories. The majority, unclassified fragments (Unknown), reflect expected plasma cell-free RNA heterogeneity, while core structural RNAs (rRNA/tRNA) remain consistent across platforms. **b** Per-sample precursor miRNA content (% of mapped reads, miRTrace). The low relative mapping rates are characteristic of total plasma-derived cell-free RNA sequencing without size selection, showing comparable target enrichment between platforms. **c** Mature miRNA library complexity. Boxplots with overlaid jitter points represent the number of distinct mature miRNAs detected per library after applying a stringent multi-sample expression filter (≥5 CPM in ≥4 samples). UG detects ∼1.85x more distinct mature miRNAs on average (median 440 vs. 236). **d** Global miRNA expression correlation. Scatter plot displaying the mean log_2_(CPM+1) expression of the 637 shared mature miRNAs passing the strict expression filter (≥5 CPM in ≥4 samples) on both platforms, averaged across all 78 samples. The red dashed line indicates the linear regression line, and the dotted line represents the y = x identity line (perfect agreement). **e** Per-sample cross-platform concordance. Distribution of Spearman’s rank correlation coefficient (ρ) and Lin’s Concordance Correlation Coefficient (CCC) calculated independently for each paired sample (n = 78) using log_2_(CPM+1) normalized expression of shared Mature miRNAs. Overlaid boxplots present the median and interquartile range, showing strong macroscopic agreement for individual library profiling. **f** Within-platform Principal Component Analysis (PCA) demonstrating biological separation. PCA was performed independently on Illumina (665 miRNAs; left) and UG (1,209 miRNAs; right) after log_2_(CPM+1) transformation and Z-score standardization. Points represent individual samples (n=78), colored by disease condition (red: PDAC; green: Control). For both platforms, the second principal component (PC2) captured statistically significant biological separation between cohorts (Illumina PC2: 5.1% variance, Welch’s t-test *P* = 1.84×10⁻⁴; UG PC2: 5.1% variance, *P* = 4.53×10⁻⁴). CPM, counts per million; CON, healthy control; PDAC, pancreatic ductal adenocarcinoma.

Among annotated reads, rRNA represented the largest fraction - Illumina ∼16% (std ∼12%) and UG ∼11% (std 8%), followed by tRNA ∼1-2%, while precursor miRNAs contributed <1% on both platforms and artifacts were negligible (Fig. 2a). Overall, annotated biotype proportions were mostly similar between platforms after decontamination, while UG exhibit a higher fraction of reads remaining unassigned to these reference biotypes.

Precursor miRNAs content (% of mapped reads) was low but consistent with what is expected from plasma cf-sRNA, averaging ∼0.66% for Illumina vs ∼0.52% for UG (std ∼0.31% vs ∼0.24%; n = 78 libraries) (Fig. 2b). The almost 25% higher relative Illumina proportion (despite lower absolute yield) suggests marginally better miRNA enrichment efficiency. Reported miRNA fractions in plasma small-RNA datasets vary widely depending on isolation strategy, library preparation, and analysis choices, ranging from low single-digit percentages depending on workflow and counting strategy to much higher proportions in others^36–38^.

### Mature miRNA expression profiles

Next, we compared mature miRNA expression between Illumina and UG. Initially, UG detected 2,032 distinct miRNAs (∼76.5% of the reference), while Illumina detected 1,308 (∼49.2%), with an overlap of 1,240 detected by both platforms. The substantially higher raw detection rate of UG reflects its greater sequencing depth, which increases statistical power to capture rare, lowly expressed transcripts. Even after removing lowly expressed reads (see Methods) UG retained 1,209 miRNAs and Illumina retained 665, with 637 shared - meaning nearly 95.8% of all Illumina passed miRNAs are also confidently detected by UG. This high concordance in the filtered set suggests that Illumina reliably captures the core, abundantly expressed miRNAs, but its lower depth might not be sufficient for detection of lower-abundance species. The ∼724 UG-exclusive miRNAs post-filter may represent biologically real, lowly expressed miRNAs that are accessible only with deeper sequencing.

To further assess this, we look at the library complexity, measured as the number of distinct miRNAs detected per sample (Fig. 2c). Library complexity was noticeably higher for UG (mean 427; std 101) than Illumina (230; std 76; n = 78 libraries). Both platforms’ counts lie within the expected complexity range for cell-free plasma^39–41^, which is highly method-dependent^42,43^. UG’s higher detection rate likely reflects its vastly higher raw sequencing depth, which causes deep sampling of low-abundance targets. General RNA-seq and small-RNA-seq methods papers show that library complexity and the number of detected features increase with sequencing depth, especially for low-abundance miRNAs near the detection limit^44,45^.

We evaluated global correlation in mean mature miRNA expression between platforms across the 637 shared, filtered miRNAs (≥5 CPM in ≥4 samples on both platforms), following log_2_(CPM+1) variance stabilization and per-miRNA mean computation across all 78 samples (Fig. 2d). The Spearman correlation is strong and significant (*r* = 0.836, *P* = 7.70×10^−168^), with high-abundance miRNAs clustering tightly around the *y = x* identity line. This indicates agreement in the high-expression regime. As seen with a previous RNA-seq comparison study^20^, at lower expression levels, scatter increases and the regression line moves upward from the identity line, reflecting a tendency for UG to report higher mean expression values for lowly expressed miRNAs. This is consistent with UG’s broader detection range, but potentially also influenced by platform-specific background noise at low abundance. Because global averaging can obscure individual library differences, we performed a paired, per-sample concordance assessment (Fig. 2e). Across the 78 individual sample pairs, the platforms demonstrated moderate concordance (mean Lin’s Concordance Correlation Coefficient (CCC) = 0.70; mean Spearman = 0.66). These values should be interpreted in the context of sparse small-RNA count data suspectable to reproducibility challenges. As an internal same-platform reference, we examined an independent small RNA-seq dataset resequencing on an Illumina machine of the same sample. Using the same mature miRNA log2(CPM+1)-based comparison strategy, this comparison yielded Spearman’s ρ = 0.69 and Lin’s CCC = 0.71 across 898 shared miRNAs (data not shown), similar to the per-sample cross-platform values observed here. Thus, the observed Illumina-UG concordance is within the range expected for sparse mature miRNA profiles and does not by itself indicate a platform-specific failure.

To measure the intensity-dependent scatter that we observed globally, we performed a Bland-Altman analysis (Fig. S8a) and divided the per-sample CCC by expression tertiles (Fig. S8b). Global Bland–Altman analysis concluded a modest positive mean difference for UG relative to Illumina. Limits of agreement (−6.85 to +8.33) were broad but centered near zero, indicating a small systematic positive bias (+0.74) for UG rather than major platform disagreement (Fig. S8a). Highly abundant mature miRNAs have strong sample-level reproducibility (mean CCC = 0.69), yet cross-platform concordance was lower within the medium (mean CCC = 0.25) and low (mean CCC = 0.21) abundance tiers (Fig. S8b). Altogether, these findings determine that while the platforms are highly interchangeable for well-expressed miRNAs, lower-abundance features require platform-aware analytical thresholds.

To validate that each platform independently captured biological variation between disease and control samples, PCA was performed separately on the Illumina (665 miRNAs) and UG (1,209 miRNAs) filtered datasets, following log2(CPM + 1) transformation and Z-score standardization. Both platforms showed biologically meaningful structure (Fig. 2f). On the Illumina dataset, PC1 explained 11.5% of variance (Fig. 2f, left panel) and did not separate conditions, while PC2 (Fig. 2f, left panel; 5.1%) demonstrated a statistically significant separation between PDAC and control samples (Welch’s t-test, *P* = 1.84×10⁻⁴). Similarly, the UG dataset showed that PC1 captured 8.6% of variance without clear condition-based clustering, while PC2 (Fig. 2f, right panel; 5.1%) exhibited separation of PDAC from control (*P* = 4.53×10⁻⁴). The similar pattern across both platforms, with the biological signal consistently emerging along PC2 rather than PC1, suggests that the primary axis is dominated by inter-individual differences unrelated to disease status, while the secondary axis captures disease-associated expression changes. The similarity of PC2-driven separation and similar p-value ranges across platforms demonstrate that both Illumina and UG independently detect the biological signal.

### Differential expression analysis

Subsequently, we performed differential expression analysis to quantify and compare the disease-associated miRNA signatures detected by each technology. Illumina (665 miRNAs) and UG (1,209 miRNAs) were analyzed separately. This per-platform analysis approach allows for the assessment of each technology’s sensitivity for detecting disease-associated signals and provides the basis for evaluating cross-platform concordance in biomarker identification.

To assess cross-platform concordance in differential expression analysis, we compared fold changes for the 637 miRNAs shared between both platforms (Fig. 3). The correlation was statistically significant and positive (Pearson *r* = 0.567, *P* < 2.2×10^−16^). This indicates that miRNAs identified as upregulated in the Illumina dataset were generally identified as upregulated in the UG dataset, and vice versa for downregulated miRNAs. Yet the moderate correlation coefficient shows that there are still meaningful differences between the two technologies, suggesting platform-specific technical factors (such as differences in adapter ligation efficiency, PCR amplification biases, or sequence-specific detection sensitivities). The scatterplot illustrates three distinct classes: first, miRNAs that reach statistical significance (*Padj* < 0.05, |*log₂FC*| > 1.0) on both platforms, which cluster tightly along the identity line and represent the strongest biomarker candidates. Second, miRNAs that are significant only on Illumina or UG (Fig. 3), which probably reflect platform-specific differences in measurement precision or within-group variability for individual miRNAs. And third, non-significant miRNAs, which show broad scatter with low effect sizes on both platforms. Among the top concordant miRNAs labeled (Fig. 3), the directionality is highly consistent across platforms, supporting the biological validity of these signals. The linear regression fit closely follows the identity line, confirming agreement in fold change directionality. Importantly, while UG’s absolute counts exhibit higher feature-level noise due to indel errors, the relative biological differences between PDAC and control groups are preserved at the level of differential expression analysis, demonstrating that the biological signal exists.

**Fig. 3:**
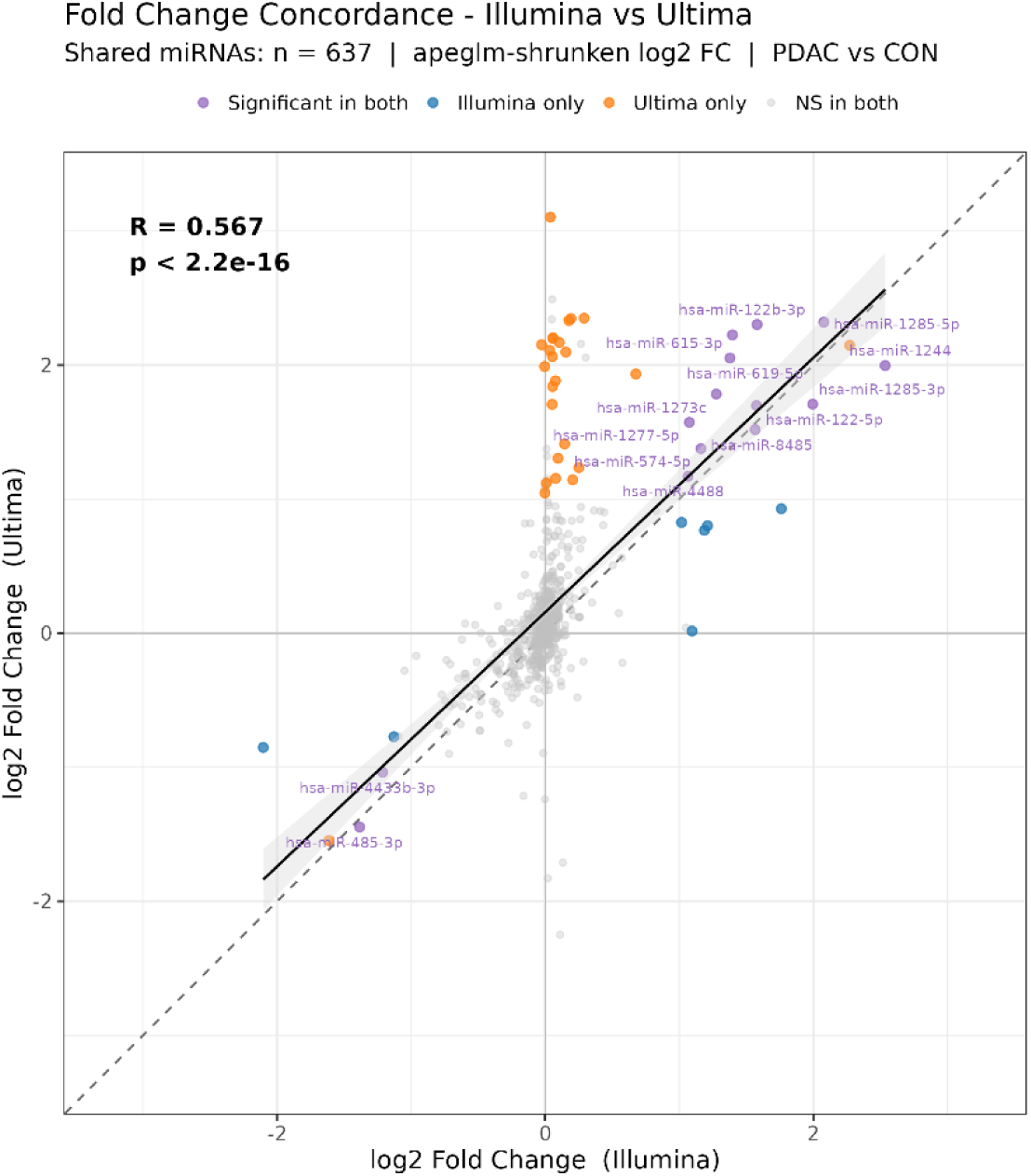
Cross-platform concordance of miRNA fold change estimates between Illumina and UG. Scatterplot comparing log₂ fold changes (PDAC vs. CON) for the 637 miRNAs shared between Illumina and UG platforms following independent DESeq2 analysis. Each point represents one miRNA, colored by: purple = significant on both platforms (*Padj* < 0.05, |log₂FC| > 1.0), blue = Illumina only, orange = UG only, grey = non-significant on both. The dashed grey line represents perfect concordance, while the black line shows the linear regression fit with 95% confidence interval (grey shading). Top concordant miRNAs (highest combined effect sizes among those significant on both platforms) are labeled. CON, healthy control; PDAC, pancreatic ductal adenocarcinoma; FC, fold change.

Using the significance threshold of *Padj* < 0.05 and |*log₂FC*| > 1.0, DESeq2 identified a total of 21 significant miRNAs in Illumina and 46 in UG (Table S4). The significant miRNAs were divided into three mutually exclusive categories: Illumina-only (n=7), UG-only (n=32), and those shared across both platforms (n=14), corresponding directly to the colored points (Fig. 3). The 14 miRNAs significant on both platforms (Fig. 3 colored purple) were entirely directionally concordant - 12 were upregulated and 2 were downregulated on both platforms. Taken together, the union of significant miRNAs across both platforms comprised 53 unique miRNAs, which were used as the complete set for disease association analysis.

As a sensitivity analysis, we compared the primary model (condition-only) with an adjusted model including age, sex, and extraction date. The adjusted model demonstrated concordance with the primary analysis. Across both platforms, the adjusted model preserved the direction of effect for all shared significant miRNAs, and correlations of shrunken log_2_ fold changes with Spearman coefficient of 0.71 and 0.70, Illumina and UG respectively (Fig. S10a and S10b). We concluded that the overall effect estimates remain similar and the adjusted specification did not alter the direction of the observed signal. As such, we retained the primary model for downstream analyses in this technical-note context.

### Disease association analysis

To study the differentially expressed miRNAs identified by each platform using existing literature/databases, we queried all significant miRNAs against two curated databases: the Human miRNA Disease Database (HMDD v4.0) and the Database of Differentially Expressed MiRNAs in human Cancers (dbDEMC 3.0). Database queries were based on two levels: first, disease classification prioritizing PDAC-specific entries, second, broader pancreatic annotations and other cancers. dbDEMC 3.0 contains study-based records of differentially expressed miRNAs across cancer types, which were subsequently filtered to pancreatic cancer in Homo sapiens. Additionally, the resource provides regulation information per study, allowing assessment of expression changes are consistent with previously published pancreatic cancer data.

Across the 53 significant miRNAs identified by DESeq2 - 7 Illumina-only, 32 UG-only, and 14 detected on both platforms - the majority had prior literature support in at least one database (Fig. S11). Fifty out of 53 miRNAs (94.3%) were present in dbDEMC 3.0 with pancreatic cancer evidence, and 15 out of 53 (28.3%) had PDAC-specific entries in HMDD v4.0. Therefore, the detected differential expression largely reflects biologically relevant signals rather than technical artifacts.

Both databases, PDAC evidence (n = 15, Fig. S11): 15 miRNAs supported by both HMDD and dbDEMC with pancreatic cancer evidence. Of these, 5 were significant on both platforms (hsa-miR-574-5p, hsa-miR-615-3p, hsa-miR-122-5p, hsa-miR-485-3p, hsa-miR-4433b-3p). The remaining 10 were platform-specific (3 Illumina, 7 UG). Among the 15 miRNAs, directional agreement with published literature was strongest for hsa-miR-574-5p (12/17 dbDEMC studies concordant, upregulated), hsa-miR-122-5p (7/11 concordant, upregulated), and hsa-miR-4433b-3p (8/9 concordant, downregulated). hsa-miR-485-3p showed a consistent down regulation across both platforms, although its direction in the literature is mixed according to dbDEMC.

dbDEMC only, pancreatic evidence (n = 35, Fig. S11): The largest group, with mostly UG-only miRNAs, had pancreatic cancer support only in dbDEMC. Several of these were significant on both platforms and showed high directional concordance with the database, including hsa-miR-619-5p (8/9 dbDEMC studies concordant, upregulated) and hsa-miR-4488 (6/6 studies concordant, upregulated). Yet, several UG-only miRNAs, such as hsa-miR-8071 and hsa-miR-6734-3p, showed complete directional discordance with dbDEMC (0 concordant studies).

HMDD other cancer evidence only (n = 3, Fig. S11): Three miRNAs significant on both platforms (hsa-miR-122b-3p, hsa-miR-1244, and hsa-miR-8485) had HMDD evidence only in general cancer contexts and no dbDEMC pancreatic cancer entries.

### Pathway analysis

Collective functional impact of the differentially expressed miRNAs was identified by each platform. For this, we performed Over-Representation Analysis (ORA) using validated miRNA-target interactions from miRTarBase (see Methods). Across the miRNA groups, ORA identified extensive pathway enrichment, with Reactome having the richest signal and KEGG showing more variable coverage (Fig. 4). As expected, the largest numbers of significant pathways were observed for the UG-significant (46 miRNAs; 25 KEGG, 7 GO, 72 Reactome) and UG-only sets (32 miRNAs; 60 KEGG, 12 GO, 108 Reactome). Yet it is important to note, the UP-intersection (12 miRNAs) still produced several enriched pathways (1 KEGG, 4 GO, 36 Reactome), indicating that the validated targets of shared and platform-specific miRNAs converge on relevant signaling.

**Fig. 4:**
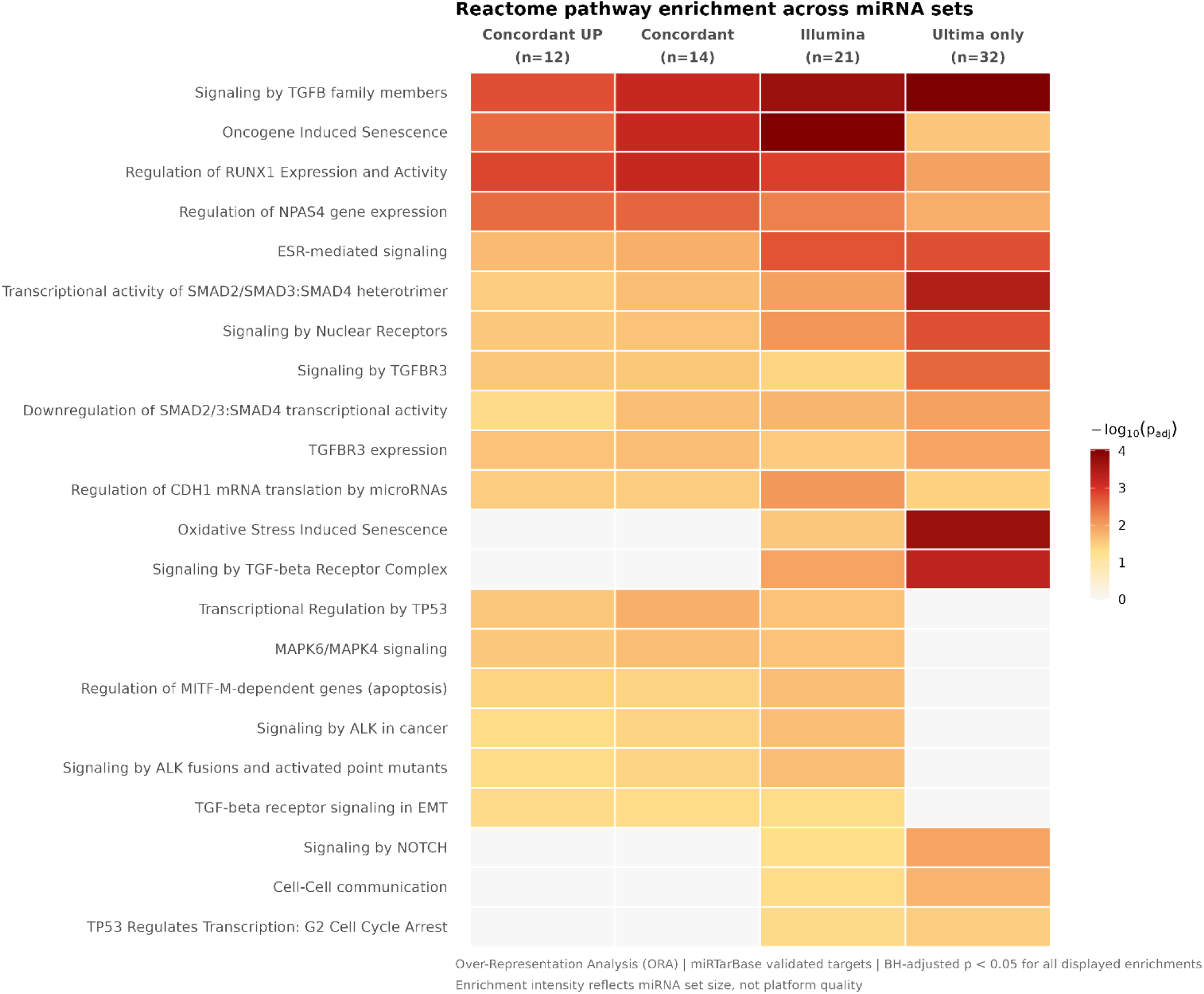
Reactome pathway enrichment across miRNA sets. Heatmap showing Over-Representation Analysis (ORA) results for 22 curated Reactome pathways across four miRNA sets derived from differential expression analysis (PDAC vs. healthy controls). Columns represent miRNA sets of increasing size: upregulated miRNAs detected concordantly by both platforms (Concordant UP, n=12), all miRNAs detected concordantly by both platforms (Concordant, n=14), all Illumina-significant miRNAs (Illumina, n=21), and UG-exclusive miRNAs (UG only, n=32). Color intensity reflects enrichment significance (−log₁₀ adjusted p-value); white cells indicate pathways not significantly enriched in that set. All pathways presented reached statistical significance (BH-adjusted p < 0.05). Pathway terms were taken from a broader ORA output to retain biologically relevant PDAC-associated processes – terms appearing in at least two miRNA sets and lacking PDAC biological relevance were excluded. ORA was performed using validated miRNA-target interactions from miRTarBase, with the union of all tested miRNAs as background.

The UP-intersection miRNA set (n = 12), representing upregulated miRNAs detected by both platforms, showed over-representation of validated targets in PDAC-relevant pathways. Reactome analysis (Fig. 4) identified 36 significant pathways (*q* < 0.2), featuring PDAC drivers such as FGFR signaling, EGFR family signaling, PDGF and VEGFR signaling, RAS/MAPK cascade activation, PI3K–AKT signaling, regulation of cell cycle progression, apoptosis execution, integrin signaling, and extracellular matrix organization. KEGG enrichment showed in EGFR tyrosine kinase inhibitor resistance as the top pathway, while GO Biological Process identified regulation of cell proliferation and positive regulation of apoptotic process. Interestingly, the UG-only set (n = 32) produced a richer functional signal (60 KEGG, 12 GO BP, 108 Reactome) than the UG-significant set (46 miRNAs; 25 KEGG, 7 GO, 72 Reactome), which included similar PDAC-relevant pathways such as RTK signaling, cell cycle regulation, apoptosis, extracellular matrix remodeling, and immune signaling, but with greater heterogeneity than the intersecting upregulated subset. This suggests that the highly abundant, shared miRNAs target a broad array of genes, diluting the pathway enrichment statistics when pooled.

Isolating the lowly expressed, UG-exclusive miRNAs (n = 32) appear to represent biologically real signals rather than platform-specific artifacts. In the fold change concordance plot (Fig. 3), UG-only hits cluster in the upper-right quadrant with positive fold changes on both platforms, but below Illumina’s significance threshold. They follow the same regression trend line (*R* = 0.567) as concordant hits. This pattern most likely indicates that UG’s deeper sequencing detects the same directional signal as Illumina, just at lower expression levels that do not quite reach statistical significance (rather than random artifacts that would scatter unpredictably across all quadrants). Most importantly, 94% of all significant miRNAs (50/53), including the platform-specific ones, had prior pancreatic cancer documentation in dbDEMC 3.0, indicating established biological relevance. In addition, ORA of UG-only targets produced pathway enrichment results (60 KEGG, 108 Reactome pathways) that generally overlap with the miRNAs in Illumina, and contain PDAC-relevant processes such as RTK signaling, RAS/MAPK activation, PI3K/AKT signaling, cell cycle regulation, and extracellular matrix remodeling. This suggests that the UG-exclusive miRNAs post-filter represent biologically real, lowly expressed miRNAs that are accessible via deeper sequencing.

## Discussion

This study is the first direct comparison of Illumina and UG sequencing for plasma-derived cf-sRNA profiling using the same PDAC and control libraries. Both platforms have a consistent disease-associated mature miRNA signal despite technical differences in sequencing output and alignment behavior. Overall, at substantially higher sequencing depth, UG provided higher mature miRNA read counts, and broader miRNA detection, while Illumina showed slightly better relative enrichment of mature miRNAs and higher post-filter genome alignment rates.

Several findings support the conclusion that both platforms are analytically suitable for plasma cf-sRNA studies - at least on the level of mature miRNA quantification and PDAC-control comparison. First, both platforms produced high-quality short-read data after filtering, both showed largely similar post-decontamination biotype compositions, and independently captured statistically significant separations between PDAC and control samples in within-platform PCA. Second, expression levels for the 637 shared filtered miRNAs were strongly correlated across platforms, especially in the higher-abundance range. This shows that the core circulating miRNA signal is strong with both sequencing technologies. Third, differentially expressed miRNAs were directionally concordant overall, and the miRNAs significant on both platforms clustered near the identity line, which supports reproducibility of the strongest disease-associated signal.

Nonetheless, the results suggest that platform choice does affect sensitivity and feature recovery in plasma cf-sRNA sequencing. UG detected more total and filtered miRNAs, and identified more differentially expressed miRNAs than Illumina, which is most likely explained by its greater sequencing depth and resulted in better sampling of low-abundance miRNAs. In contrast, Illumina had a higher proportion of mature miRNA reads relative to total short cfRNA input and higher genome alignment rates. This is consistent with a cleaner enrichment profile and known differences between substitution-dominant SBS chemistry and the higher indel rate reported for UG chemistry, which may be especially relevant for very short RNA fragments.

An important point is that the additional signals detected by UG do not appear to be dominated by random technical noise. Although many significant miRNAs were platform-specific, the UG-only signals usually followed the same fold-change direction as Illumina. They lay along the same global regression trend and were often supported by past pancreatic cancer evidence in dbDEMC and HMDD. In addition, targets of the UG-only miRNAs resulted in enrichment for pathways already connected to PDAC biology, including receptor tyrosine kinase signaling, RAS/MAPK and PI3K/AKT signaling, cell-cycle regulation, apoptosis, and extracellular matrix remodeling. Thus, these observations are more consistent with higher sensitivity to lower-abundance biologically relevant miRNAs than with only artifacts, although some platform-specific calls likely still are a result of chemistry- or workflow-dependent bias. It should be noted that HMDD v4.0 and dbDEMC 3.0 are populated mostly by studies using Illumina sequencing or hybridization-based platforms and database-based corroboration is not entirely platform-neutral. In addition, the pathway inferences rely on literature-curated targets and lack PDAC-context orthogonal validation (e.g., AGO2-CLIP or perturbation assays). Thus, while these observations are consistent with higher sensitivity to lower-abundance biologically relevant miRNAs on the UG platform, orthogonal quantification is required to definitively resolve the UG-exclusive signal.

The study also presents the greater technical challenges of plasma cf-sRNA profiling. In both datasets, mature miRNAs represented only a very small fraction of total short cfRNA reads. Substantial percentages of reads were removed as cfDNA/cDNA, ncRNA, and rRNA contamination, and a large proportion of genome-mapped reads remained unassigned to the standard annotated biotypes. This fits the fragmented, heterogeneous composition of extracellular RNA in plasma and helps explain why small differences in sequencing chemistry or mapping behavior can continue to platform effects visible in downstream analyses. This is also consistent with the PERMANOVA result (Tables S3a and S3b) that showed that platform explained more variance than disease status, even though disease-associated separation was still detected within each platform independently.

In addition to canonical transcripts, plasma cfRNA contains microRNA isoforms (isomiRs). These are biological sequence variants that occur due to post-transcriptional modifications such as 5′ or 3′ trimming and nucleotide additions. Our supplementary analysis of these variants further demonstrates the impact of platform-specific technical biases. Both platforms showed the expected dominance of 3′ end variants and a strong 18-19nt signal for specific circulating miRNAs but were different in the ratio between trimming and addition subtypes (Figure S7).

This suggests that end-variant calling is sensitive to platform-specific error modes. Because the main differential expression analysis was performed at the parent miRNA locus level, these differences are not likely to have significantly changed the conclusions of the study, but they suggest that isomiR-focused analyses may require caution when comparing between technologies.

This study has several limitations. First, it was conducted in a single case-control cohort and evaluated one low-input plasma small-RNA workflow, one library-preparation strategy, and one analytical framework centered on mature miRNAs. Accordingly, the findings may not generalize directly to other cohorts, pre-analytical conditions, library chemistries, or analyses focused on other small-RNA classes or isomiRs. Second, although sequencing the same biological libraries on both platforms is a major strength because it minimizes biological variability, the libraries were originally prepared in an Illumina-compatible format and subsequently adapted for UG sequencing rather than generated with platform-native protocols. Therefore, some of the observed differences may reflect library-platform compatibility in addition to sequencing chemistry itself. Third, the platforms were not depth-matched, and UG produced substantially more reads; as a result, its higher miRNA detection rate and greater number of differentially expressed miRNAs likely reflect a combination of deeper sampling and platform-specific performance, rather than chemistry alone. Finally, no orthogonal validation or independent external cohort was used to confirm the shared or platform-specific differentially expressed miRNAs. Thus, the present work should be interpreted primarily as a technical benchmarking study, rather than as a definitive biomarker discovery study. Orthogonal assays (such as RT-qPCR or ddPCR) and external replication are required before these signatures can be considered for clinical validation.

To conclude, we demonstrate that UG sequencing, similar to Illumina, is a viable platform for plasma cf-sRNA sequencing and may offer a practical advantage when broader miRNA detection and greater sensitivity to low-abundance signals are desired. Illumina has advantages in relative mature miRNA enrichment and alignment efficiency. Most importantly, the core PDAC-associated biological signal was preserved across both platforms, and the most correlated miRNAs and pathway-level results were consistent with prior literature. Future studies should test whether these conclusions hold in larger cohorts, across additional plasma and extracellular RNA workflows, and should validate whether the extra miRNAs recovered by deeper sequencing improve classification, early detection, or monitoring performance in PDAC liquid biopsy settings.

## Methods

### Cohort aggregation and sample acquisition

A cohort of 78 viable samples was assembled. This cohort comprised 39 control subjects and 39 PDAC patients. The sample collection spanned several months, from May 2023 to January 2024, and was conducted across two medical institutions in Israel: (i) Hadassah Medical Center (Jerusalem, Israel), the study protocol was reviewed and approved by their Institutional Review Board (IRB) (Approval number: HMO-0198-14); (ii) Sheba Medical Center (Tel Hashomer, Israel), the study protocol was reviewed and approved by their IRB (Approval number: SMC-9534-22). Written informed consent was obtained from all participants prior to sample collection. Post-collection and plasma isolation, the samples were preserved at −80 °C and dispatched to our laboratory for subsequent processing. Each sample was accompanied by demographic data, including age, BMI and gender. Samples from collection through processing were recorded on the Gotsho LIMS system (www.gotsho.com, accessed 2 August 2024).

### Cell-free small RNA isolation from blood plasma

For the purification of cell-free nucleic acids from plasma samples, we used the QI-Aamp Circulating Nucleic Acid Kit (Qiagen, Venlo, The Netherlands)^46^, which provides a structured method for this purpose. The process begins with the lysis of plasma samples using proteinase K and Buffer ACL under denaturing conditions to release nucleic acids from their protein or vesicle-bound states. After lysis, Buffer ACB is added to adjust the binding conditions, allowing the nucleic acids to adhere to a silica membrane within the QIAamp Mini columns. The bound nucleic acids are then washed with Buffer ACW1, Buffer ACW2, and ethanol to remove residual contaminants. Finally, the nucleic acids are eluted in Buffer AVE. This method is designed to handle large sample volumes and facilitate the recovery of nucleic acids, making it applicable for the sequencing of small cell-free RNA fragments. The samples’ output was assessed using random sampling using Qubit RNA High Sensitivity (Invitrogen, Carlsbad, CA, USA)^47^ and Bioanalyzer RNA Small RNA chip (Agilent, Santa Clara, CA, USA)^48^ to ensure minimal loss of material.

### Library preparation for cell-free small RNA-seq

For the purpose of turning the purified cell-free nucleic acids into a library suitable for sequencing, we used the SMARTer smRNA-Seq Kit for Illumina (Takara, Shiga, Japan)^49^. This kit is specifically designed for the preparation of sequencing libraries from small RNA fragments. The process involves polyadenylation of the RNA, followed by cDNA synthesis using template switching technology, which incorporates adapters required for sequencing. After amplification and size selection to ensure the correct library size, the samples are prepared for sequencing. The library preparation was carried out using standardized protocols suitable for the sequencing of small cell-free RNA fragments. The cDNA libraries prepared were assessed with Bioanalyzer DNA High Sensitivity chip (Agilent, Santa Clara, CA, USA).

Subsequently, these same libraries were further modified by adding UG-specific adapters so that the same biological libraries could be sequenced on both platforms (Illumina NovaSeq 6000 and Ultima UG 100). Due to minimal initial plasma volumes, no size selection and additional RNA purification steps were performed.

### Small RNA-seq quality control, stepwise alignment and miRNA quantification

Raw FASTQ files from both platforms underwent initial quality control trimming adapters and polyA, 5’ bias, polyG/X tails, low-quality bases (Q≥20), and length filtering (18-40 nt), yielding total short cfRNA reads for downstream analysis.

To isolate mature miRNAs from total short nucleic acids (including rRNA/tRNA fragments and cfDNA contaminants inherent to low-input plasma extraction), the reads underwent decontamination using sequential Bowtie2 mapping to custom indices: rRNA, piRNA, tRNA (GtRNAdb), cDNA (Ensembl GRCh38), and ncRNA (excluding miRNA loci). This step discards matches ≥10 nt, retaining unaligned reads for subsequent steps.

We aligned the remaining reads to the reference miRNA database using Bowtie against mature miRNA and precursor (hairpin) loci. The resulting alignments are processed by SAMtools for quantification and miRTrace provides miRNA precursor loci (hairpins) quality control metrics, including the proportion of reads assigned to precursor miRNAs versus other RNA classes and sequencing artifacts.

We further aligned reads to a mature miRNA reference FASTA (miRBase v22), containing 2,656 distinct annotated human mature miRNA sequences, then filtered the raw count matrices: miRNAs required ≥5 CPM (total library-normalized counts) in ≥4 samples. UG retained 1,209 miRNAs and Illumina retained 665, with 637 shared. Quality metrics, RNA biotype proportions, and gene-level counts were compared using Python.

For paired, per-sample cross-platform concordance analysis, normalized expression vectors [log2(CPM+1)] for the shared mature miRNAs were compared independently for each of the 78 paired samples. Within each pair, we calculated Spearman’s rank correlation coefficient (ρ) and Lin’s Concordance Correlation Coefficient (CCC). To quantify absolute systematic differences and define 95% limits of agreement, we performed a global Bland–Altman analysis using all paired miRNA observations. Finally, the shared mature miRNAs were stratified into equal tertiles (Low, Medium, and High abundance) based on their global mean CPM. Per-sample Lin’s CCC was then re-calculated iteratively within each abundance tier.

### Principal Component Analysis

To assess the main transcriptomic variance and evaluate platform batch effects, PCA was performed on the log_2_(CPM+1) normalized expression values of the shared mature miRNAs. Features were standardized (Z-score transformed) before dimensionality reduction. Global PCA was performed on the combined Illumina and UG datasets to test platform separation, while intra-platform PCAs were generated independently for each technology to check biological stratification (PDAC vs. Control). We then performed unadjusted, single-factor Permutational Multivariate Analysis of Variance (PERMANOVA), and multi-factor PERMANOVA to make sure that the concordant biological separation observed on both platforms was not driven by pre-analytical batch effects. To adjust for potential confounders, we applied a sequential multi-way PERMANOVA (vegan::adonis2 in R), structured to remove out variance attributed to clinical site and collection month, and then evaluate sequencing platform and biological condition. Statistical significance was assessed using 9,999 permutations, and approximate R^2^ values were calculated to quantify the proportion of variance explained by each factor.

### IsomiR and mirtop analysis

mirtop (v0.5.1) processed miRBase-aligned mature reads to annotate canonical/isomiR variants, quantifying modifications (add/del/sub at 5’/3’/internal) and generating stats. This enables objective assessment of isomiR diversity, important for cf-sRNA where post-transcriptional variants may carry biomarker value.

### Differential expression analysis and count normalization

Differential expression analysis was performed using DESeq2 in R. The statistical design checked the contrast between PDAC and control samples (design = ∼ label), with PDAC as the reference level, such that positive *log₂FC* indicates higher expression in PDAC relative to controls. To improve effect size estimation and reduce noise for low-count features, we applied log fold change shrinkage using the apeglm method. Differentially expressed miRNAs were defined using standard thresholds of adjusted p-value (*Padj*) < 0.05 (Benjamini-Hochberg correction) and absolute shrunken *log₂FC* > 1.0. As a sensitivity check, we also ran an adjusted model incorporating age, sex, and extraction date, and calculated concordance with the primary model by comparing shrunken log_2_ fold changes across all tested miRNAs.

### Disease association analysis

We queried the DESeq2-significant miRNAs identified across the two sequencing platforms against two curated literature resources: the Human miRNA Disease Database (HMDD v4.0) and the Database of Differentially Expressed MiRNAs in human Cancers (dbDEMC 3.0). HMDD was used to assess previous experimentally validated miRNA-disease associations, we prioritized PDAC-specific entries and then broader pancreatic or other cancer annotations. dbDEMC was restricted to human pancreatic cancer records and used to evaluate whether the observed direction of differential expression matched previously reported UP/DOWN changes.

### Pathway analysis

We performed Over-Representation Analysis (ORA) using validated miRNA-target interactions from miRTarBase using the multiMiR R package. Only experimentally supported interactions were retained, including both strong evidence interactions (Functional MTI like luciferase reporter assays, western blot) and weaker but still experimental interactions (Functional MTI Weak like PAR-CLIP, microarray). The aim is to provide a high-confidence set of miRNA-target gene relationships. Target retrieval was performed against the union of all miRNAs tested across both platforms (not just the significant subset) so that we have a miRNA target gene set as the background for enrichment testing, rather than the background of all human genes in general. ORA was then applied to seven miRNA sets: all Illumina-significant miRNAs (n=21), all UG-significant miRNAs (n=46), the intersection (n = 14), its upregulated and downregulated subsets (UP, n=12; DOWN, n=2), and the platform-specific Illumina-only (n=7) and UG-only (n=32) groups. For each group, the target genes were tested for over-representation using KEGG, GO Biological Process, and Reactome pathway databases using clusterProfiler and ReactomePA, with a BH-adjusted p-value threshold of 0.05 and q-value cutoff of 0.2.

## Acknowledgements

We thank members of the Shomron laboratory for handling the biological processing and DNA extraction along valuable support in data acquisition and technical consultation. We thank the Sheba Pancreatic Cancer Center and Hadassah oncology department for providing cancer and control patients cohort. This study was supported in part by a fellowship from the Edmond J. Safra Center for Bioinformatics at Tel-Aviv University.

## Author contributions statement

A.L. performed the bioinformatic and statistical analysis and written the final manuscript. H.V. conceived and planned the study, provided guidance and revised final manuscript. R.S. conducted the biological experiments and N.S. provided overall scientific supervision, critically revised the manuscript, read and approved the final version.

## Data availability statement

The raw sequencing data generated in this study has been deposited in the European Genome-phenome Archive (EGA) under the study accession number EGAD50000002771.

