## supplementary information for "Comparative analysis of Illumina and Ultima-Genomics sequencing for plasma cell-free small RNA profiling in pancreatic cancer"

#### Fig. S1: Absolute read count distributions for Total Short cfRNA and Mature miRNA datasets.

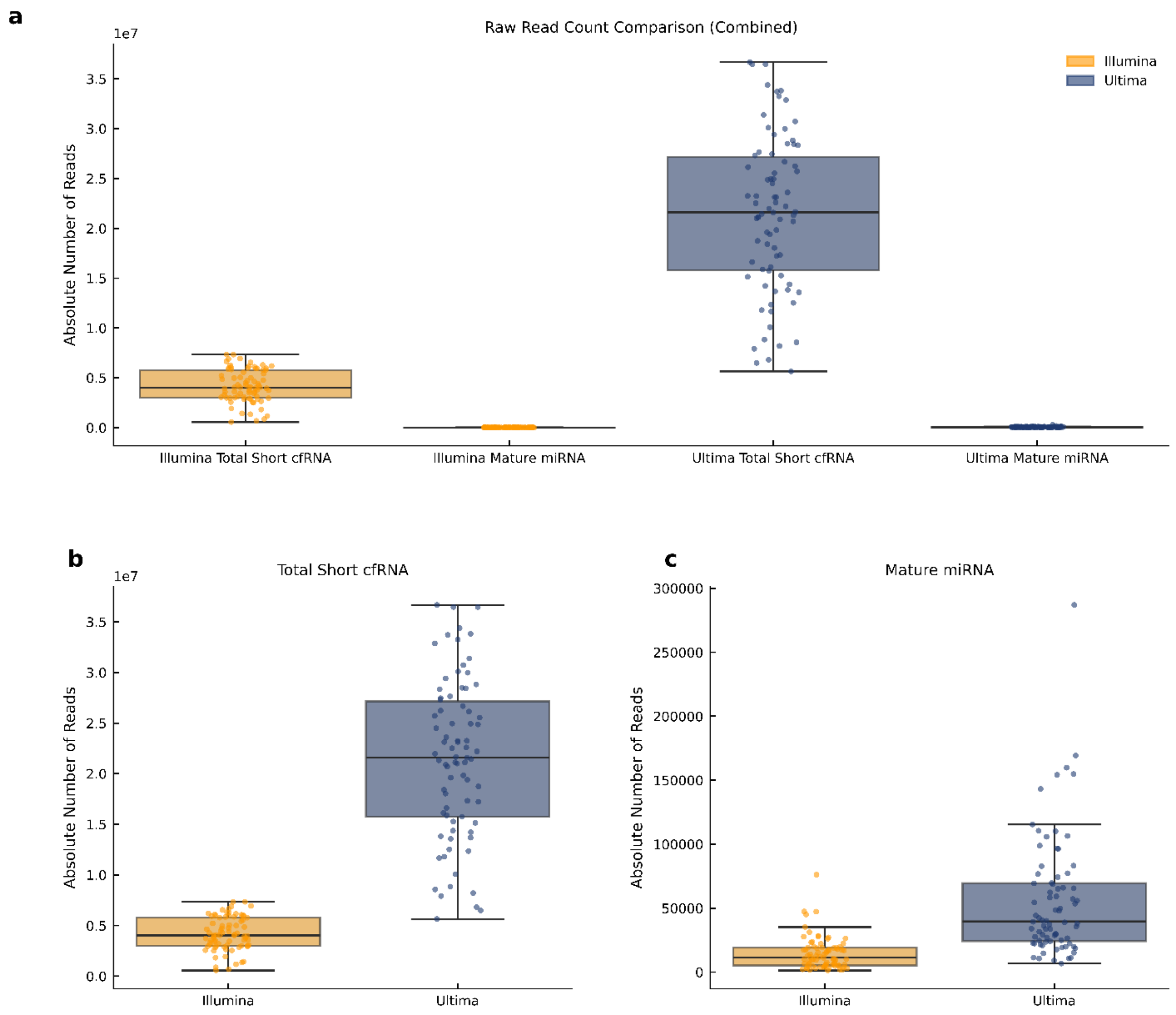


**a** Per-library read counts for total short cfRNA (post-trimming/QC), and after mature miRNA alignment, for both platforms. Boxplots with jitter points show medians, interquartile ranges, and individual libraries (n = 78). **b** Comparing only Illumina vs UG reads counts for total short cfRNA (post-trimming/QC). **c** Comparison of Illumina and UG read counts after mature miRNA alignment.

**Table S1: Summary of sequence read yields and mature miRNA recovery across sequencing platforms.**

| Platform | Samples | Metric | N | Mean | SD | Min | 25th Pctl | Median | 75th Pctl | Max |
|---|---|---|---|---|---|---|---|---|---|---|
| Illumina | Total Short cfRNA | Raw Reads | 78 | 4,193,562 | 1,646,419 | 559,947 | 3,007,391 | 4,028,798 | 5,773,712 | 7,338,135 |
| | | % Total Yield | 78 | 1.28 | 0.50 | 0.17 | 0.92 | 1.23 | 1.77 | 2.24 |
| | Mature miRNA | Raw Reads | 78 | 14,592 | 12,677 | 1,281 | 5,190 | 11,413 | 19,015 | 76,131 |
| | | % Total Yield | 78 | 0.0045 | 0.0039 | 0.0004 | 0.0016 | 0.0035 | 0.0058 | 0.0233 |
| UG | Total Short cfRNA | Raw Reads | 78 | 21,480,356 | 7,816,102 | 5,647,640 | 15,775,045 | 21,611,743 | 27,152,811 | 36,675,784 |
| | | % Total Yield | 78 | 1.28 | 0.47 | 0.34 | 0.94 | 1.29 | 1.62 | 2.19 |
| | Mature miRNA | Raw Reads | 78 | 55,980 | 47,037 | 6,677 | 24,169 | 39,668 | 69,585 | 287,009 |
| | | % Total Yield | 78 | 0.0033 | 0.0028 | 0.0004 | 0.0014 | 0.0024 | 0.0042 | 0.0171 |

Read count statistics are shown for 78 identically processed plasma samples sequenced on both Illumina NovaSeq 6000 and Ultima-Genomics UG100 platforms. Total Short cfRNA (Raw Reads) indicates the absolute number of reads retained per sample after strict adapter trimming and length filtering. To account for the platforms' differing overall capacities, % Total Yield expresses each sample's read count as a percentage of the entire sequencing run's usable depth (a balanced 78-plex pool would average 1.28% per sample). Mature miRNA (Raw Reads) represents the absolute number of reads successfully aligning to the miRBase mature database following decontamination. Mature miRNA (% Total Yield) represents the final normalized proportion of the total sequencing run dedicated to biologically valid mature miRNA sequences for each sample.

#### Contamination filtering

Contamination filtering removed ~55-60% of total short cfRNA reads across platforms, with cDNA dominating - Illumina mean 37% (std ~10%) and UG 34% (std ~9%). This was followed by ncRNA, averaging ~13% (std ~5%) and 12% (std ~4%) for Illumina and UG respectively. Afterwards, rRNA with an average of ~11% (std ~8%) for Illumina and ~9% (std ~7%) for UG. Remaining unaligned reads (target-enriched for miRNAs) comprised ~39% (~11% std; Illumina) vs ~46% (~9% std; UG), reflecting UG's higher initial yield. piRNA/tRNA contributions were negligible (<0.3% for both).

Across libraries, a substantial fraction of total short cfRNA reads passed contamination filtering, with UG retaining a higher proportion than Illumina - mean ~46% vs ~39% and median ~44% vs ~38% (~9% std vs ~11% std). Despite this shift, both platforms showed broad yet overlapping distributions (IQR roughly ~31-47% for Illumina and ~39-52% for UG), indicating similar overall contamination burdens with modestly greater usable yield from UG.

**Fig. S2: Contamination filtering composition and sequence retention rates.**

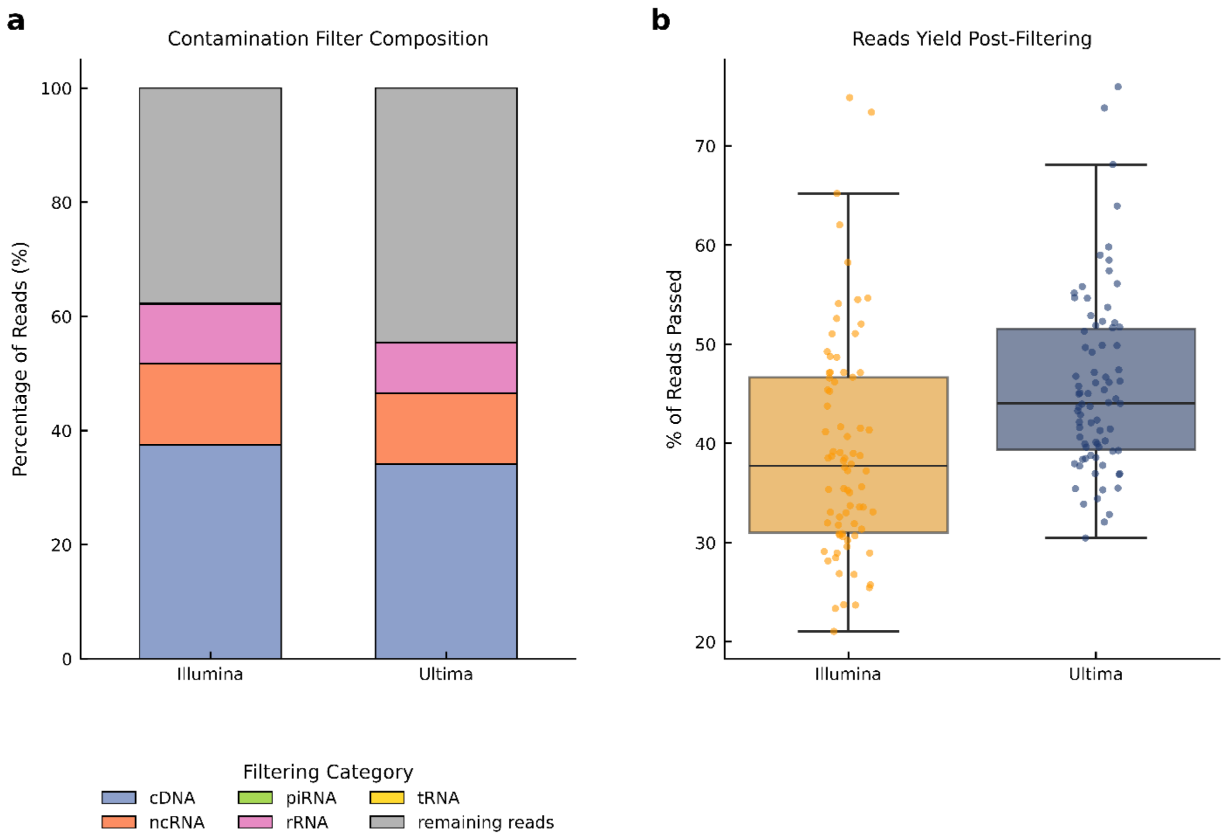


**a** Mean composition of reads removed during contamination filtering by biotype. Stacked bars show proportions (n=78 libraries per platform); cDNA/ncRNA/rRNA dominate losses in both platforms, validating consistent decontamination effectiveness. **b** Percentage of total short cfRNA reads retained after sequential contamination filtering. Boxplots with jitter display per library percentages (n = 78 per platform); UG retains slightly more reads on average, but variability overlaps across platforms.

**Table S2: Stepwise read counts during contamination filtering. Mean reads and percentage of total post-adapter-trimmed input reads removed at each sequential filtering stage (rRNA, tRNA, piRNA, cDNA, ncRNA) and passed to mature miRNA alignment. Reported per platform across all samples.**

| Filtering Stage / Biotype | Illumina [Mean Reads] | Illumina [% of Total] | UG [Mean Reads] | UG [% of Total] |
|---|---|---|---|---|
| **Total Input Reads (Post-Adapter Trimmed)** | 4,193,562 | 100.00 | 21,480,356 | 100.00 |
| **rRNA** | 433,472 | 10.34 | 1,899,634 | 8.84 |
| **tRNA** | 9,919 | 0.24 | 29,416 | 0.14 |
| **piRNA** | 755 | 0.02 | 5,775 | 0.03 |
| **cDNA** | 1,572,755 | 37.50 | 7,314,983 | 34.05 |
| **ncRNA** | 597,263 | 14.24 | 2,668,577 | 12.42 |
| **Passed to miRNA Mapping** | 1,579,398 | 37.66 | 9,561,971 | 44.51 |

The stepwise read count table shows that cDNA was the dominant filtered fraction for both platforms (Illumina: 37.5%; UG: 34.1%), followed by ncRNA (Illumina: 14.2%; UG: 12.4%) and rRNA (Illumina: 10.3%; UG: 8.8%). This is consistent with the contamination composition shown in Fig. S2a and with the longer-fragment signal in Fig. 1e. Following all filtering steps, 37.7% of Illumina reads and 44.5% of UG reads were passed to mature miRNA alignment. This once again shows UG's greater retention of mappable small RNA content. This difference may contribute to its higher miRNA detection yield downstream.

**Fig. S3: Locus-specific read length distributions highlighting native circulating short isoforms.**

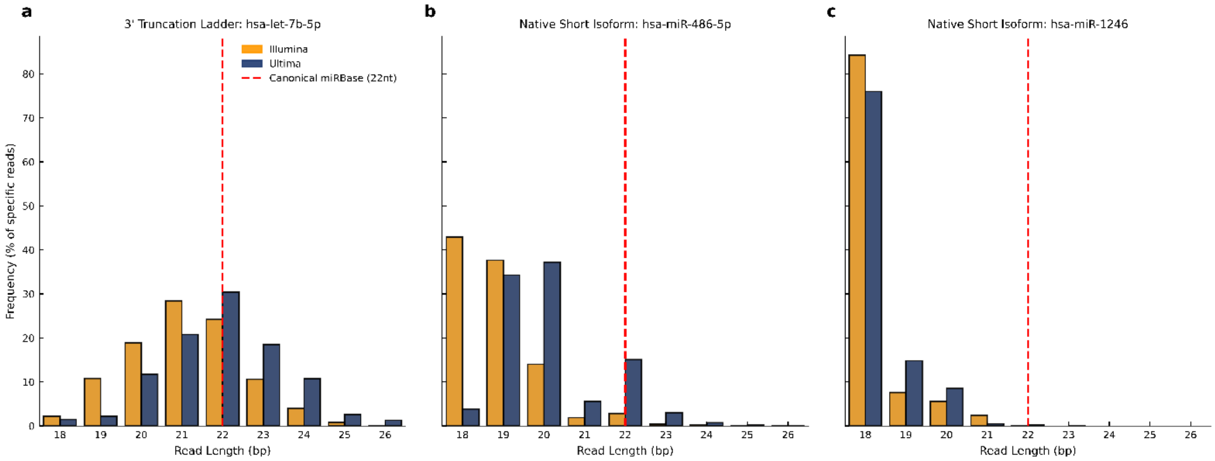


Locus-specific read length distributions reveal the 18-19nt peak in plasma small RNA sequencing. Read length frequencies (expressed as a percentage of total locus-specific mapped reads) are shown for three abundant circulating RNAs across the Illumina (orange) and UG (blue) platforms. The canonical 22nt length is indicated by the red dashed line. **a** hsa-let-7b-5p demonstrates a classic truncation ladder, with reads degrading stepwise from the canonical 22nt down to 18nt. **b** hsa-miR-486-5p natively exists in plasma as a highly stable 18-19nt isomiR, with few reads mapping to the canonical 22nt length. **c** hsa-miR-1246 similarly presents a predominant 18-19nt peak.

**Fig. S4: Per-base sequence quality profiles across sequencing platforms.**

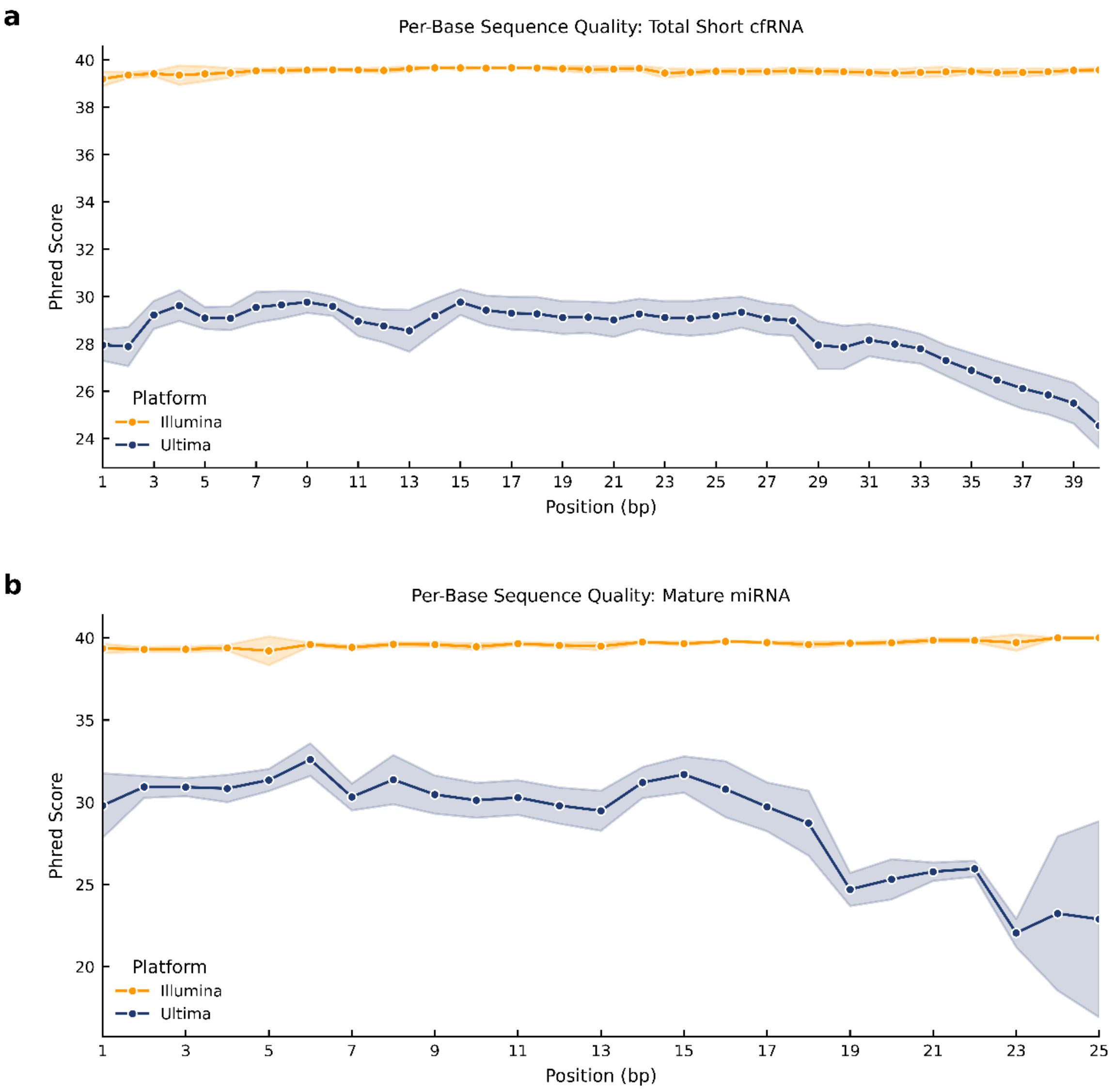


**a** Mean per-base sequence quality (Phred scores) post-adapter trimming and **b** after mature miRNA alignment. Lines depict platform averages across n=78 libraries (positions 1-25/40 bp); both platforms deliver high quality, with mature miRNA data showing higher quality overall.

Per-base quality scores (Phred) remained excellent across platforms post-adapter trimming (means 28-39 for both), with Illumina maintaining flatter profiles all the way and UG showing sharper 3' decline typical of flow chemistry. Mature miRNA reads exhibited mostly higher scores, confirming effective enrichment of high-confidence target sequences after processing.

**Fig. S5: Per-sequence GC content distribution of mature miRNAs.**

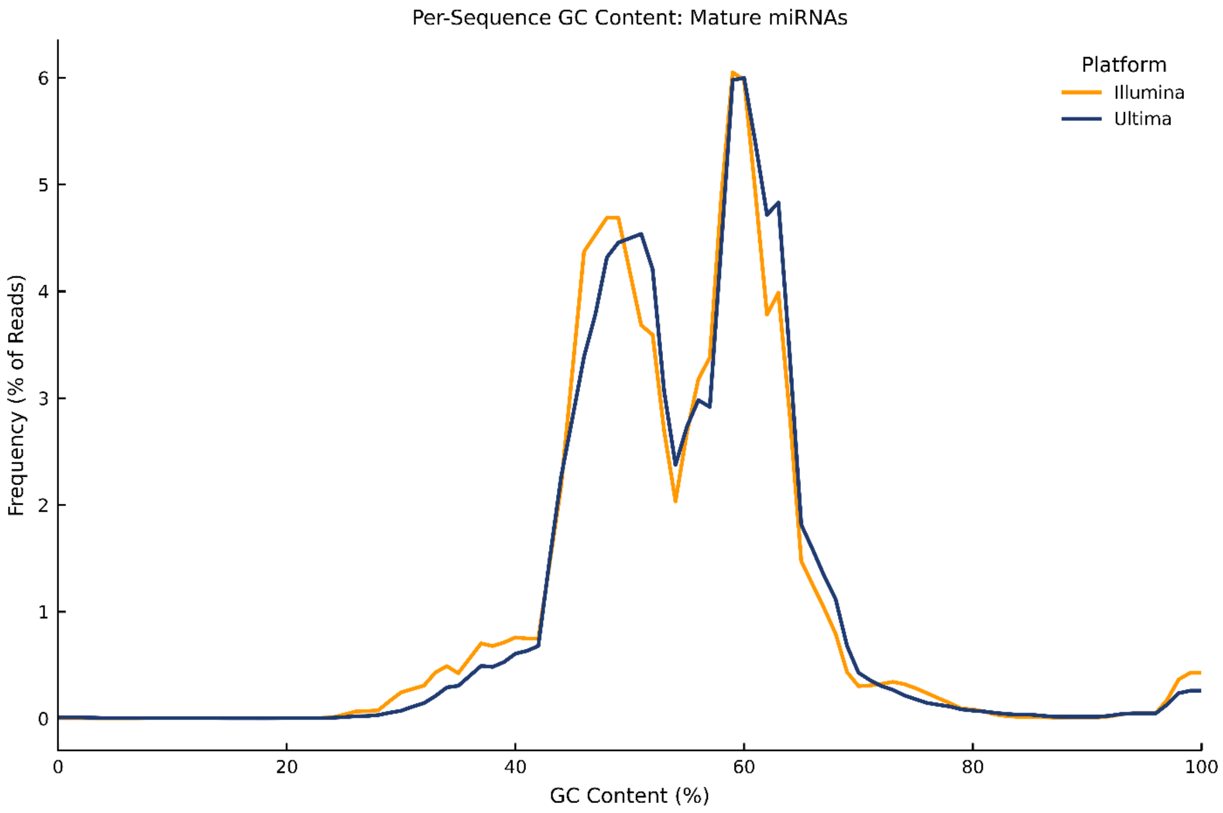


Frequency distribution of per-sequence GC content (%) in mature miRNA reads. Curves represent platform averages across n=78 libraries (0-100% GC bins); profiles are highly concordant, indicating equivalent miRNome representation.

Mature miRNA per-sequence GC content distributions were comparable between platforms, both exhibiting expected bimodal profiles with peaks at ~50% and ~60% GC, reflecting miRBase composition (means ~45-50% GC; n=78 libraries). Minor peak shifts (UG slightly right-shifted) suggest negligible platform-specific GC bias post-enrichment and filtering. The near-identical profiles across the full range indicate insignificant platform-specific GC bias post-enrichment and filtering which supports cross-platform differential expression comparisons (as GC-dependent count distortions would not disproportionately affect either platform).

**Fig. S6: Sequence duplication rates within mature miRNA libraries.**

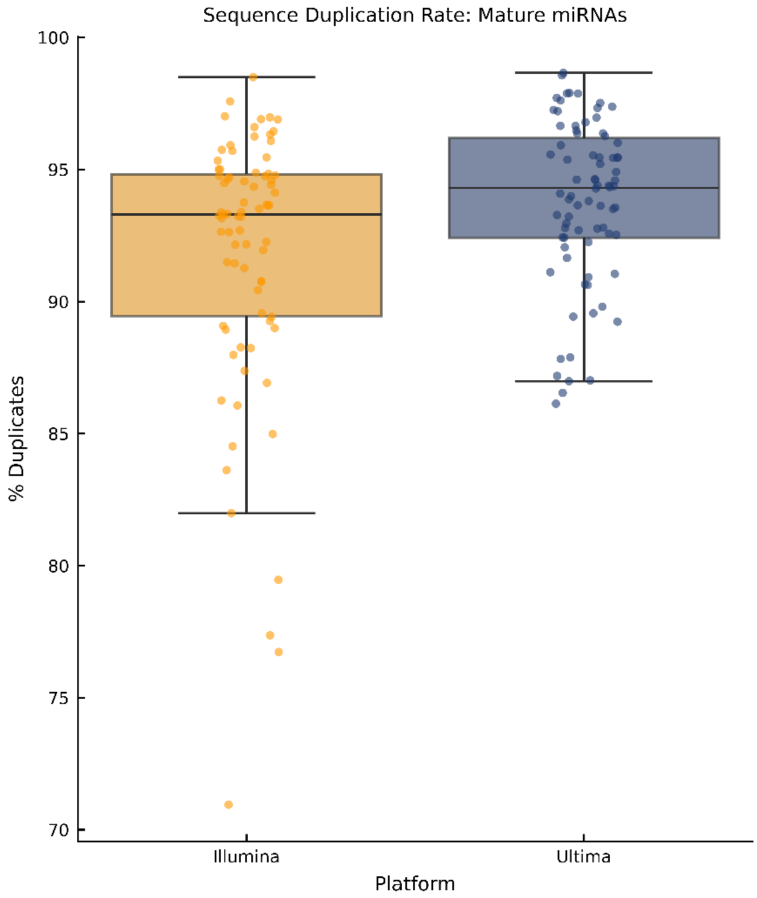


Sequence duplication rates (% duplicated reads) in mature miRNA datasets by platform. Boxplots with jitter show medians ~93-94%, interquartile ranges, and individual libraries (n=78); comparable complexity confirms equivalent library diversity post-processing.

Sequence duplication rates in mature miRNA reads were high but comparable across platforms, reflecting low-diversity cf-sRNA libraries (Illumina mean 92%; UG 94%; n=78 libraries each), with both showing relatively tight distributions (std 3-5%). The slightly higher (but overlapping) UG values align with its greater depth amplifying PCR/low-complexity duplicates. In the context of small RNA-seq, high duplication is expected given the limited numbers of mature miRNAs and the dominance of a small number of highly abundant species in plasma. Future differential expression analysis operates on raw counts without deduplication, and the consistent duplication profiles across both platforms and samples indicate that count inflation is unlikely to introduce substantial systematic bias into cross-platform comparisons.

### IsomiR and mirtop Analysis

#### Fig. S7: Comprehensive profiling of miRNA isoform (isomiR) composition and sequence diversity.

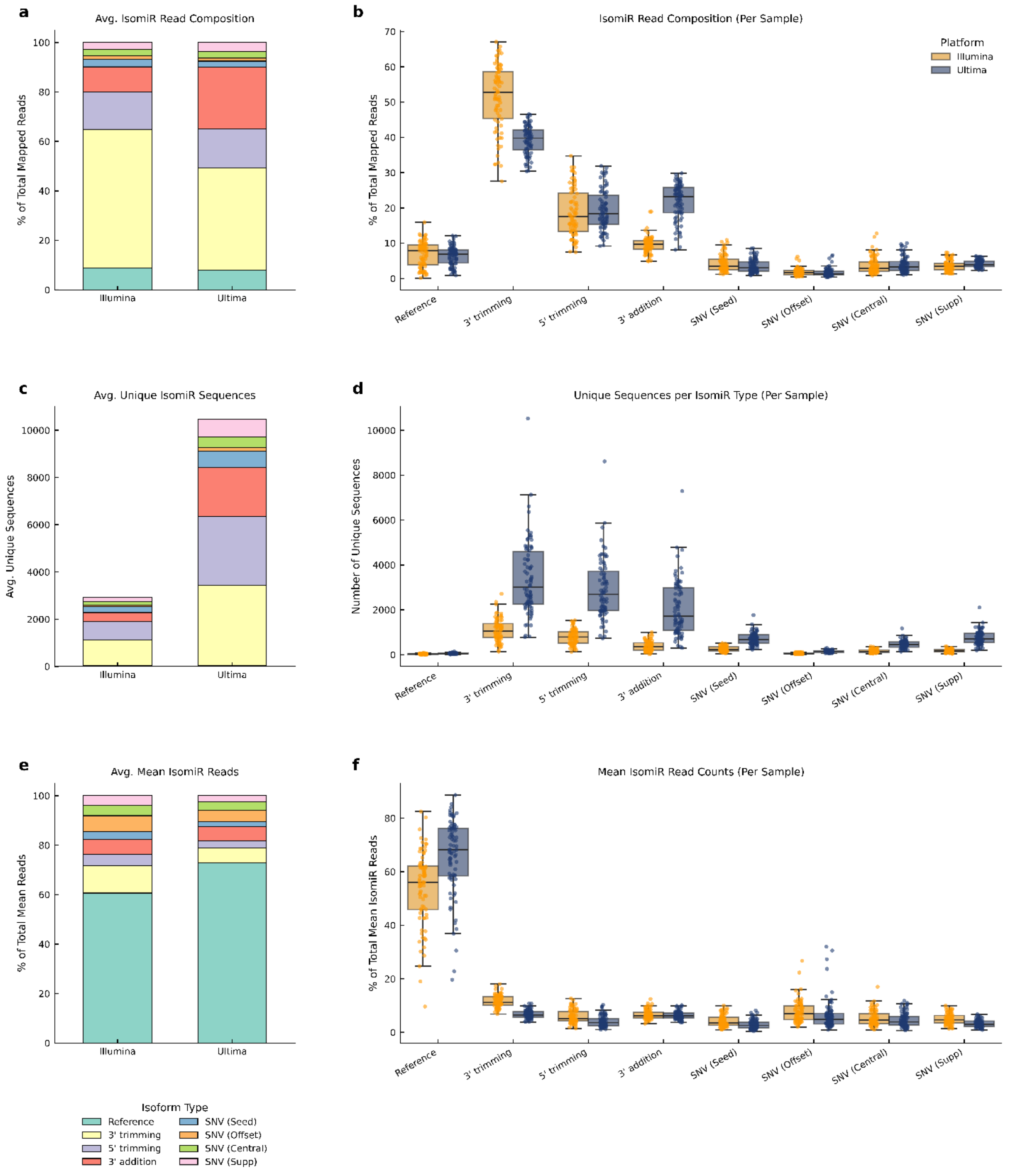


**a** mirtop isomiR subtype composition (% miRNA reads) stacked means (n=78/platform) and **b** per-library boxplots. 3' variants are most dominant. Illumina skews toward trimming over additions. **c** Unique isomiR sequences detected (total across miRNAs) stacked subtype means (n=78/platform) and **d** per-library boxplots. UG's depth yields markedly greater diversity per subtype. **e** Mean isomiR read composition per miRNA stacked subtype averages (n=78/platform) and **f** per-miRNA boxplots. Reference sequences dominate per-miRNA reads; modification subtypes are individually sparse despite their collective sequence diversity.

mirtop quantified eight canonical isomiR subtypes from miRNA-aligned reads (Fig. S7a and S7b; n=78 libraries/platform). There is cross-platform concordance of dominant 3' modifications (trimming + additions: Illumina 61%; UG 61%) and 5' trimming (19% both), with minor SNVs (~13% total) and reference types (~7%). Illumina exhibited a shifted balance within 3' variants - higher trimming (51% vs 39%; Welch t-test p<10^-20) but lower additions (10% vs 22%; Welch t-test p<10^-20). Overall, these profiles are consistent with plasma and extracellular RNA benchmarks, where 3′ length variants and 3′ additions are the dominant isomiR classes[1–3]. Thus, both platforms show the expected dominance of 3′-end isomiR heterogeneity in plasma, but they differ in the balance of 3′ trimming versus 3′ addition calls. This may reflect platform-specific differences in error modes and/or end-variant detection rather than biological differences, given that the underlying libraries are matched. This is unlikely to affect gene-level miRNA quantification or classical differential expression analyses, which collapse isomiRs to canonical miRNAs.

Aggregated across all miRNAs, UG libraries have ~2-3x more distinct sequences per isomiR subtype than Illumina (Fig. S7c and S7d; for example, 3' trimming: UG mean 3385 vs 1085; 3' addition: 2069 vs 385). Per-library boxplots confirm consistent scaling, while stacked averages highlight UG's expanded range (total unique ~11k vs 3k).

Mean read counts per miRNA (averaged over all detected miRNAs) show that reference sequences account for the majority of reads per individual miRNA (Fig. S7e and S7f; Illumina 54%, UG 66%), confirming that canonical transcripts drive bulk expression. Subtypes such as 3' trimming contribute only 7-12% of reads per miRNA on average, consistent with modification variants being individually low abundance. This is despite forming the largest pool of unique sequences collectively (Fig. S7c and S7d).

**Fig. S8: Cross-platform intensity bias and abundance-stratified concordance.**

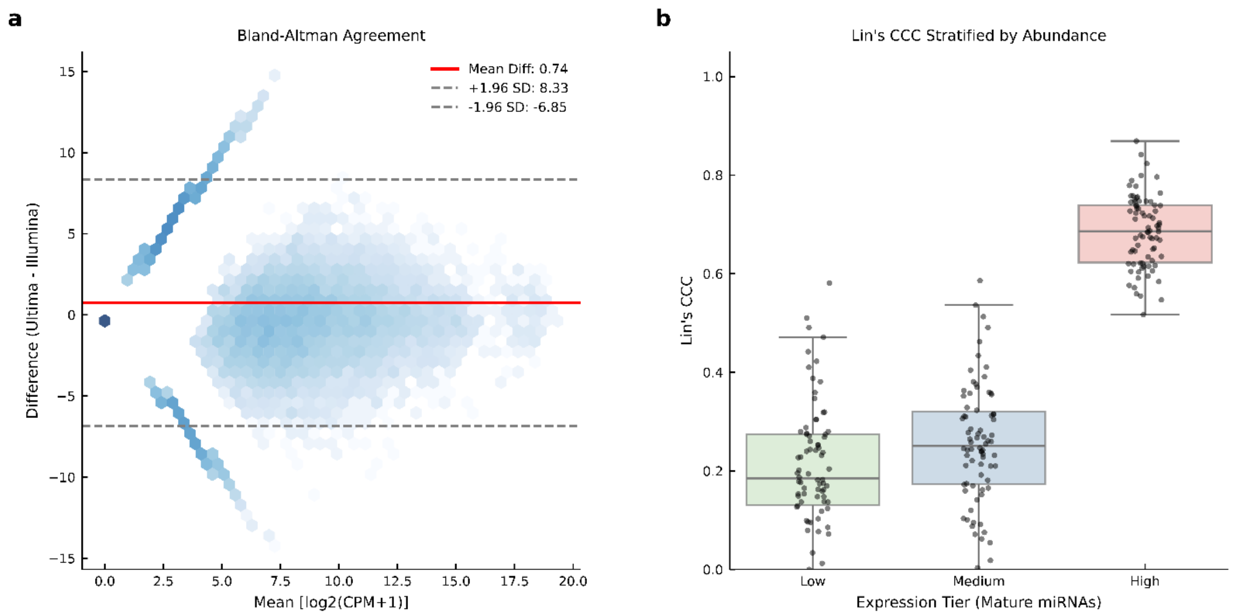


**a** Bland-Altman agreement plot comparing Illumina and UG Mature miRNA quantifications across all 78 paired libraries. The x-axis represents the mean log2(CPM+1) expression between platforms, and the y-axis displays the paired difference (UG - Illumina). Data density is represented by logarithmic hexagonal binning. The solid red line indicates the mean systematic difference and dashed grey lines the 95% limits of agreement (±1.96 SD). **b** Per-sample Lin's Concordance Correlation Coefficient (CCC) stratified by Mature miRNA abundance. Shared Mature miRNAs were divided into equal tertiles (Low, Medium, and High) based on global mean CPM. Boxplots overlay the per-sample distributions, showing robust cross-platform agreement for highly expressed features and divergence for low-abundance targets.

**Fig. S9: Combined principal component analysis evaluating cross-platform variance.**

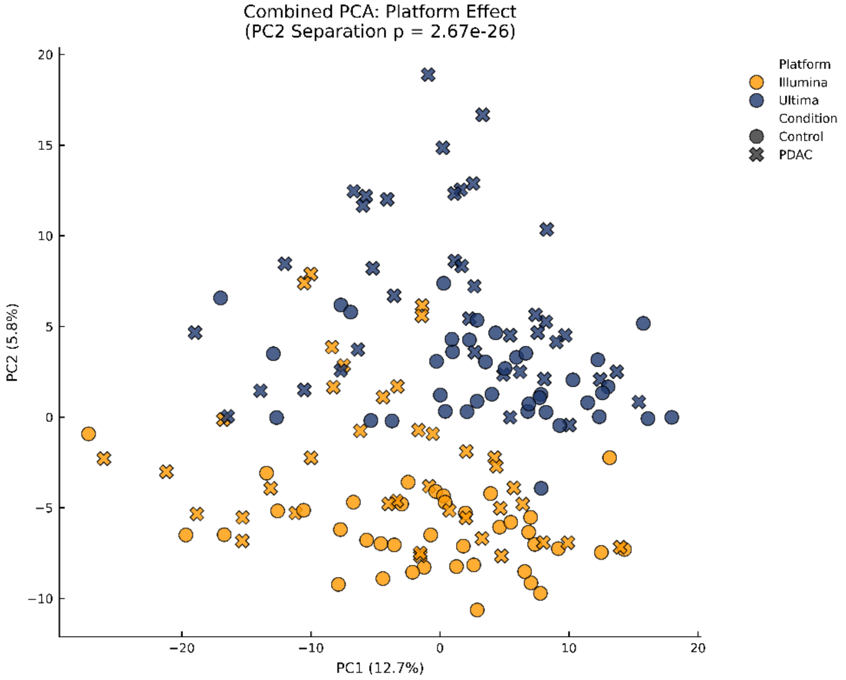


Combined PCA of Illumina and UG miRNA expression profiles. Principal component analysis (PCA) of 156 samples (78 Illumina, 78 UG) performed on the 637 shared, filtered miRNAs following log2(CPM + 1) variance stabilization and Z-score standardization. Each point represents one sample, colored by platform (orange: Illumina; blue: UG) and shaped by disease condition (circle: CON; cross: PDAC). Percentage of variance explained by each principal component is indicated on the respective axis. Platform separation along PC2 was assessed using a two-sample Welch's t-test; p-value is indicated in the title. CPM, counts per million; CON, healthy control; PDAC, pancreatic ductal adenocarcinoma.

To assess the global structure of miRNA expression across both platforms simultaneously, we performed PCA on the combined matrix of 156 samples (78 Illumina + 78 UG) using the 637 shared, filtered miRNAs, after $\log_2$(CPM+1) transformation and Z-score standardization. In the resulting plot PC2 (5.8%) shows a clear platform-level separation, with Illumina samples clustering at negative values and UG at positive values. It is highly statistically significant (Welch's t-test on PC2 scores, $p = 2.67 \times 10^{-26}$). PC1 (12.7% of variance) does not cleanly separate the two platforms. The dominant source of global expression variability reflects individual biological heterogeneity, shared by both technologies, rather than a technical effect.

**Table S3a: Single-Factor Permutational Multivariate Analysis of Variance (PERMANOVA) of global miRNA expression -** Independent, one-way PERMANOVA evaluating the variance in $log_2$(CPM+1) normalized mature miRNA expression explained by technical, pre-analytical, and biological factors individually. Significance was assessed using 9,999 permutations.

| Factor | Pseudo-*F* | *P*-value | $R^2$ |
|---|---|---|---|
| Platform (Illumina vs. Ultima) | 8.30 | 0.0001 | 0.051 |
| Condition (PDAC vs. Control) | 2.83 | 0.0001 | 0.018 |

**Table S3b: Sequential Multi-Way PERMANOVA adjusting for pre-analytical batch effects -** A sequential multi-factor PERMANOVA model aimed to evaluate the biological variance (Condition) after removing variance attributed to pre-analytical and technical covariates. The factors were evaluated in the exact sequence listed in the table (Distance ~ Month + Site + Platform + Condition). Significance was assessed using 9,999 permutations.

| Factor | df | Sum of Squares | Pseudo-*F* | *P*-value | $R^2$ |
|---|---|---|---|---|---|
| **Month** | 2 | 2091 | 1.75 | 0.0012 | 0.021 |
| **Site** | 1 | 1195 | 2 | 0.0023 | 0.012 |
| **Platform** | 1 | 5084 | 8.51 | 1.00E-04 | 0.051 |
| **Condition** | 1 | 1417 | 2.37 | 0.001 | 0.014 |
| **Residual** | 150 | 89585 | | | 0.902 |
| **Total** | 155 | 99372 | | | 1 |

To quantify the contributions of platform and disease condition to the overall expression structure, we applied a Permutational Multivariate Analysis of Variance (PERMANOVA). Platform identity was a significant factor of expression variability, accounting for approximately 5.1% of total variance ($P = 0.0001$; Table S3a). Disease condition (Control vs. PDAC) was also statistically significant, but it explained a smaller proportion of variance (~1.8%, $P = 0.0001$; Table S3a). This is about one-third of the platform effect and explains why Control and PDAC samples do not visually separate in a joint PCA plot, where PC2 separates by platform (5.8%, Fig. S9). The biological signal exists but it is outweighed by inter-individual noise and the platform-driven technical component. Cases and controls were assembled across two medical centers over an extended collection time, warranting confirmation that the biological signal was not a technical artifact of pre-analytical batch effects. We extended our analysis using a sequential multi-way PERMANOVA, structured to regress out variance attributed to collection month and clinical site, before evaluating the sequencing platform and biological condition. Importantly, even after adjusting for these potential confounders, the disease condition remained highly significant, independently explaining 1.4% of the variance ($P$ = 0.0010; Table S3b), concluding that the cross-platform disease separation reflects true tumor biology rather than a shared technical reproduction of sample-handling imbalances.

**Table S4: Summary of significant differentially expressed genes (DEGs) and shared directionality across platforms Illumina and Ultima-Genomics (UG)**

<table>
<tr><th>Platform</th><th>Total Detected miRNAs</th><th>Total DEGs [padj<0.05 & |log2FC|>1.0]</th><th>Upregulated</th><th>Downregulated</th><th>Shared DEGs</th><th>Shared Upregulated</th><th>Shared Downregulated</th></tr>
<tr><td>Illumina</td><td>665</td><td>21</td><td>17</td><td>4</td><td rowspan="2">14</td><td rowspan="2">12</td><td rowspan="2">2</td></tr>
<tr><td>UG</td><td>1209</td><td>46</td><td>44</td><td>2</td></tr>
</table>

**Fig. S10: Concordance of differential expression estimates between the primary and adjusted models.**

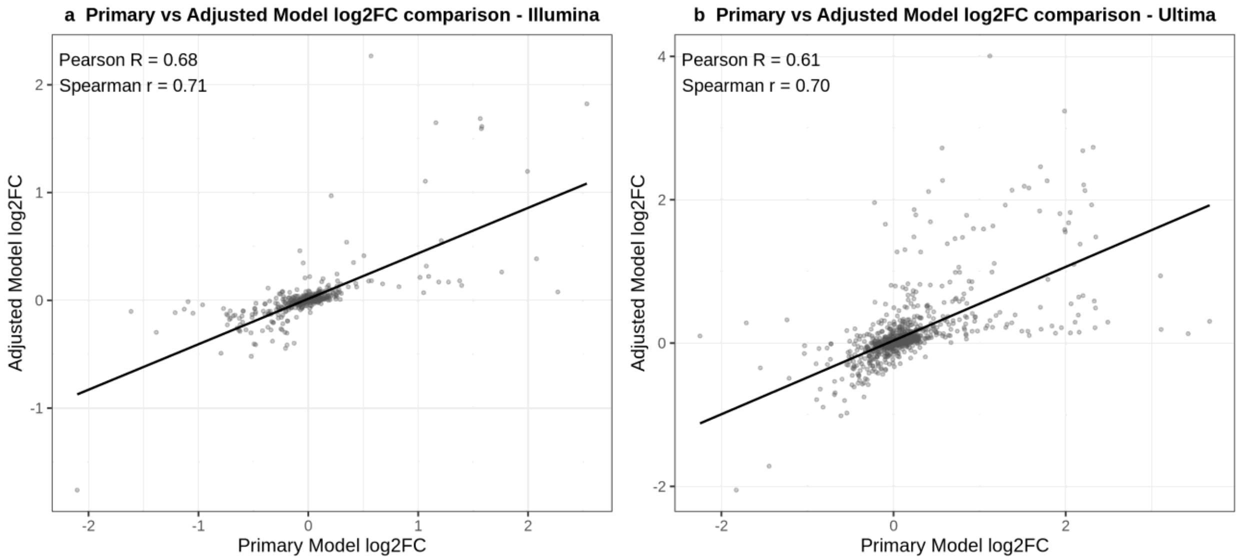


Scatterplots compare shrunken $\log_2$ fold changes from the primary ~ label (PDAC vs Control) model and the adjusted ~ age + sex + extraction date + label model for Illumina (panel a) and UG (panel b). Each point represents one miRNA; the fitted line shows the overall linear trend. Pearson and Spearman correlation coefficients are shown within each panel to summarize agreement between models.

**Fig. S11: Literature-based evidence for all differentially expressed miRNAs identified by DESeq2 (PDAC vs. healthy controls).**

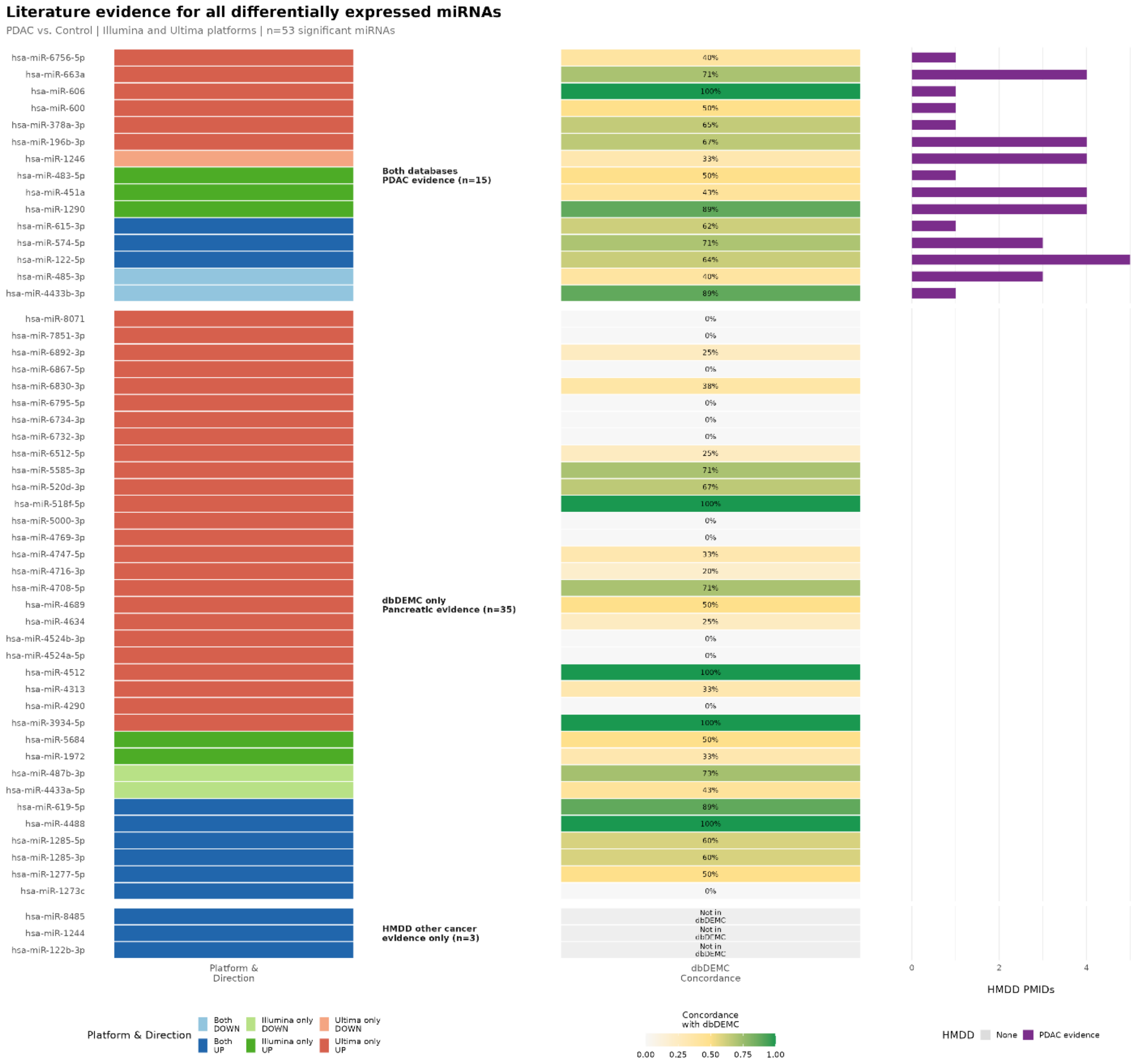


Each row represents one of the 53 significant miRNAs detected across both platforms. Rows are grouped by evidence category (as defined in the main text). Both databases, PDAC evidence (n=15): miRNAs with experimentally validated entries in both HMDD v4.0 (PDAC-specific or broader pancreatic annotations) and dbDEMC 3.0 (pancreatic cancer, Homo sapiens). dbDEMC only, pancreatic evidence (n=35): miRNAs with pancreatic cancer records in dbDEMC 3.0 only. HMDD other cancer evidence only (n=3): miRNAs with HMDD entries in general cancer contexts but no dbDEMC pancreatic cancer records. **Left panel:** Platform of detection and direction of differential expression

(UP/DOWN, PDAC vs. controls). **Middle panel:** Directional correlation with published pancreatic cancer data from dbDEMC 3.0, expressed as the percentage of dbDEMC studies reporting the same direction of expression change as observed here. Grey cells indicate no pancreatic cancer records in dbDEMC 3.0 for that miRNA. **Right panel:** Number of PubMed-indexed publications (PMIDs) supporting miRNA–disease associations in HMDD v4.0. For the Both databases, PDAC evidence group, PMIDs reflect PDAC-specific or pancreatic entries. For the HMDD other cancer evidence only group, PMIDs reflect general cancer associations.
